# Antibodies Raised Against an Aβ Oligomer Mimic Recognize Pathological Features in Alzheimer’s Disease and Associated Amyloid-Disease Brain Tissue

**DOI:** 10.1101/2023.05.11.540404

**Authors:** Adam G. Kreutzer, Chelsea Marie T. Parrocha, Sepehr Haerianardakani, Gretchen Guaglianone, Jennifer T. Nguyen, Michelle N. Diab, William Yong, Mari Perez-Rosendahl, Elizabeth Head, James S. Nowick

## Abstract

Antibodies that target the β-amyloid peptide (Aβ) and its associated assemblies are important tools in Alzheimer’s disease research and have emerged as promising Alzheimer’s disease therapies. This paper reports the creation and characterization of a triangular Aβ trimer mimic composed of Aβ_l7-36_ β-hairpins, and the generation and study of polyclonal antibodies raised against the Aβ trimer mimic. The Aβ trimer mimic is covalently stabilized by three disulfide bonds at the corners of the triangular trimer to create a homogeneous oligomer. Structural, biophysical, and cell-based studies demonstrate that the Aβ trimer mimic shares characteristics with oligomers of full-length Aβ: X-ray crystallography elucidates the high-resolution structure of the trimer and reveals that four copies of the trimer assemble to form a dodecamer; SDS-PAGE, size exclusion chromatography, and dynamic light scattering reveal that the trimer also forms higher-order assemblies in solution; cell-based toxicity assays show that the trimer elicits LDH release, decreases ATP levels, and activates caspase-3/7 mediated apoptosis. Tmmunostaining studies on brain slices from people who lived with Alzheimer’s disease as well as people who lived with Down syndrome reveal that the polyclonal antibodies raised against the Aβ trimer mimic recognize pathological features including different types of Aβ plaques and cerebral amyloid angiopathy. These findings suggest that the triangular trimer structural motif is important in Alzheimer’s disease and may thus constitute a new structurally defined molecular target for diagnostic and therapy development.

**SYNOPSIS:** A structurally defined Aβ oligomer mimic is created and studied, and antibodies raised against the Aβ oligomer mimic are used to investigate its relevance to Alzheimer’s disease.

## INTRODUCTION

Antibodies are important tools for probing biomolecular species in cells and in tissues. Antibodies are especially valuable, because of their strong affinity and excellent selectivity for peptides and proteins, as well as their ability to be used in highly sensitive fluorescent and luminescent technologies that can identify miniscule quantities of peptides and proteins. Antibodies can also provide insights into the structures and conformations of proteins in cells and in tissues.^1,2,3^ Antibodies that target monomeric, oligomeric, and fibrillar forms of the β-amyloid peptide (Aβ) are valuable tools for Alzheimer’s disease research and have emerged as potential Alzheimer’s disease therapies.^4,5,6,7,8,9,10^

In Alzheimer’s disease, the Aβ peptide self-assembles to form oligomers and fibrils. Aβ oligomers appear to be important in the pathogenesis and progression of Alzheimer’s disease, ^11,12,13,14,15,16,17,18,19,20,21,22,23,24,25,26,27,28,29^ with Aβ dimers, trimers, hexamers, and dodecamers as well as larger oligomers identified in Alzheimer’s disease brain tissue. ^30,31,32,33,34,35,36,37^ Understanding the structures of Aβ oligomers and Aβ fibrils is crucial for understanding the molecular basis of Alzheimer’s disease and should lead to better diagnostics and therapies for Alzheimer’s disease. The structures of different Aβ *fibril* polymorphs have begun to emerge, owing to advances in cryo-EM and solid-state NMR spectroscopy.^38,39,40,41,42,43,44,45,46,47,48^ In spite of the tremendous advances in amyloid structural biology, the structures of Aβ *oligomers* remain largely unknown.^49^ High-resolution structural elucidation of Aβ oligomers by X-ray crystallography, NMR spectroscopy, or cryo-EM is hindered by challenges in preparing stable, homogeneous Aβ oligomers *in vitro* or isolating sufficient quantities of stable, homogeneous biogenic Aβ oligomers from tissue. These same challenges have also hindered the generation of antibodies against homogeneous structurally defined Aβ oligomers.

The diversity of aggregates that Aβ forms has inspired several approaches for generating Aβ antibodies as tools and probes for identifying Aβ and its many aggregates *in vitro* and in the brain. The 6E10 and 4G8 monoclonal antibodies-among the most extensively used Aβ antibodies in Alzheimer’s disease research-were generated by immunizing mice with a peptide fragment that encompassed the *N*-terminal half of Aβ (Aβ_1-24_).^50,51^ The A11 and OC polyclonal antibodies-among the first “conformation-dependent” Aβ antibodies that distinguished Aβ oligomers and Aβ fibrils-were generated by immunizing rabbits with Aβ_40_ oligomers (A11) or Aβ_42_ fibrils (OC) prepared *in vitro*.^52,53,54^ These conformation-dependent antibodies have allowed researchers to probe the structures of Aβ oligomers as well as Aβ fibrils in mouse and human brain tissues and fluids.^55,56,57,58,59,60,61^ The 1C22 monoclonal antibody-an Aβ antibody that preferentially recognizes Aβ aggregates and not Aβ monomers-was generated by immunizing mice with a disulfide-crosslinked dimer of an Aβ_40_ variant with cysteine in place of Ser_26_.^62,63,64^ The ACU193 monoclonal antibody-an Aβ antibody that is highly selective for specific types Aβ oligomers-was generated by immunizing mice with Aβ-derived diffusible ligands (ADDLs), a type of Aβ oligomer prepared by aggregating full-length Aβ *in vitro*.^65^ Hundreds of other Aβ antibodies have been raised against various forms of Aβ including Aβ peptide fragments, Aβ oligomers, and Aβ fibrils prepared under different *in vitro* conditions, and Aβ isolated from Alzheimer’s disease brains.^66^

The Aβ antigens used to generate Aβ antibodies selective for aggregated forms of Aβ contain a mixture of oligomers or fibrils with inherently diverse epitopes and undefined molecular structures. While antibodies raised against these mixtures can distinguish different aggregation states of Aβ, the lack of high-resolution structural characterization of the Aβ antigens precludes structural correlation of the *in vitro*-prepared oligomers or fibrils with oligomers or fibrils in the brain. Antibodies raised against structurally defined Aβ oligomers, with known high-resolution structures, may help shed light on the structures of the Aβ oligomers that form in the brain or serve as potential immunotherapies for Alzheimer’s disease.

This paper reports the generation and study of antibodies raised against a homogeneous structurally defined triangular trimer derived from Aβ. We first detail the design, synthesis, and X-ray crystallographic structure of the triangular trimer, and demonstrate through a series of biophysical and cell-based experiments that the triangular trimer shares many characteristics with oligomers of full-length Aβ. We then describe the generation and study of polyclonal antibodies raised against the triangular trimer. To our knowledge, these are the first antibodies raised against an Aβ-derived oligomer with a known high-resolution structure. We use these antibodies to investigate the relationship between the triangular trimer and Aβ assemblies in post-mortem brain tissue from people who lived with Alzheimer’s disease and Down syndrome, as well as brain tissue from 5xFAD transgenic mice.

## RESULTS AND DISCUSSION

### Design and Synthesis of the Covalently Stabilized Triangular Trimer 2AT-L

β-Hairpins have emerged as important structural motifs adopted by the Aβ peptide in both the oligomeric and fibrillar state.^67,68,69,70^ β-Hairpins are the simplest type of β-sheet, comprising two antiparallel hydrogen-bonded β-strands connected by a loop. Several Aβ β-hairpins have been described in which the central and *C*-terminal regions of the Aβ peptide comprise the β-strands of the β-hairpin.^71,72,73^ In one example, Hard *et al*. elucidated the NMR structure of an Aβ_17-36_ β-hairpin bound to an affibody.^74^ In subsequent studies, Hard *et al*. covalently stabilized Aβ_40_ and Aβ_42_ in a β-hairpin conformation by installing a cross-strand intramolecular disulfide bond, and demonstrated that these stabilized Aβ β-hairpins assemble to form soluble oligomers that recapitulate many characteristics of Aβ oligomers.^75,76^

To gain insights into the high-resolution structures of Aβ oligomers, our laboratory has pioneered macrocyclic β-hairpin peptides that mimic Aβ β-hairpins.^77,78^ These β-hairpin peptides contain chemical modifications that stabilize the peptides in a β-hairpin conformation and limit their propensity to aggregate. These modifications enable crystallization and elucidation of the X-ray crystallographic structures of the oligomers that the peptides can form. Using this approach, we have discovered that β-hairpin peptides that mimic Aβ_l7-36_ β-hairpins assemble to form triangular trimers that further assemble to form higher-order oligomers, such as hexamers and dodecamers.

We designed peptide 2AM-L to mimic an Aβ_l7-36_ β-hairpin (Figures lA-C). 2AM-L contains a 8-linked ornithine turn unit that connects the *N*- and *C*-termini of the peptide and helps enforce a β-hairpin conformation. To improve solubility of the peptide and prevent uncontrolled aggregation, 2AM-L also contains an *N*-methyl group on the amide backbone of Phe_20_, and the charged isostere of methionine, ornithine, at position 35. Previous X-ray crystallographic studies of three closely related peptide analogues of 2AM-L revealed that these peptides assemble to form triangular trimers (Figures Sl). While these 2AM-L analogues assemble to form triangular trimers at the high concentrations of X-ray crystallography (>l mM), these analogues and 2AM-L do not appear to form a triangular trimer at low, more biologically meaningful concentrations (<50 µM). For this reason, covalent stabilization of the triangular trimer is needed to study its structural, biophysical, and biological properties.^79,80^ Covalent stabilization of the triangular trimer also ensures oligomer homogeneity by eliminating the monomer-oligomer equilibrium that would occur for monomers that assemble to form trimers or other oligomers.

**Figure 1.**
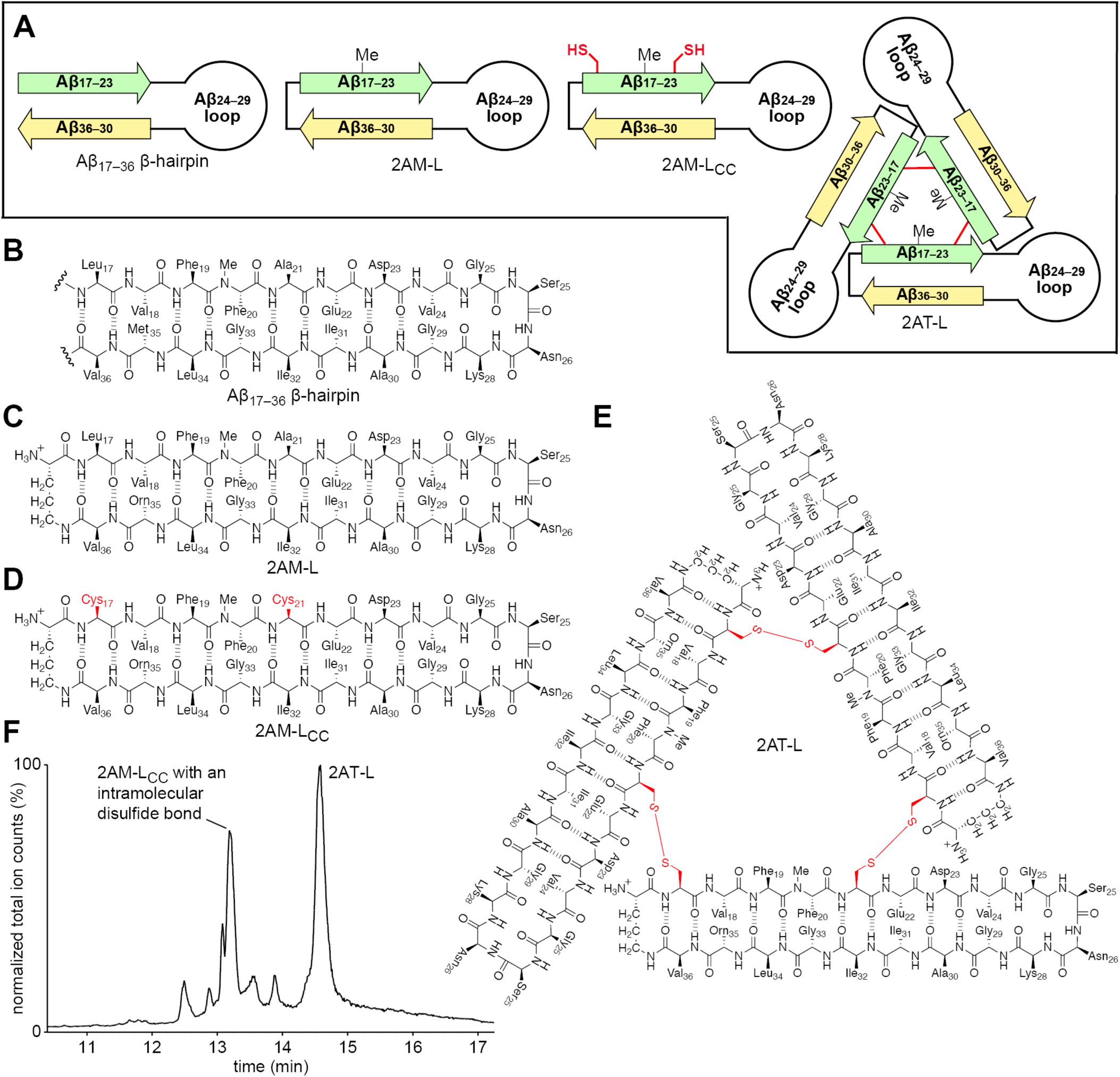
Design and synthesis of the covalently stabilized triangular trimer 2AT-L. ***(A)*** Cartoons illustrating the design of 2AM-L, 2AM-L_CC_, and 2AT-L and their relationship to an Aβ_l7-36_ β-hairpin. ***(β-E)*** Chemical structures of an Aβ_l7-36_ β-hairpin, 2AM-L, 2AM-L_CC_, and 2AT-L. ***(F)*** LC-MS trace of the oxidation reaction mixture of 2AM-L_CC_ to form 2AT-L after 48 hours in 20% DMSO with triethylamine. The two major products that form during the oxidation reaction are indicated on the trace-the desired species 2AT-L, and 2AM-L_CC_ that contains an intramolecular disulfide bond.

We designed 2AT-L as a covalently stabilized analogue of a triangular trimer formed by 2AM-L (Figures 1A and E). The design of 2AT-L is based on the previously reported X-ray crystallographic structures of triangular trimers composed of β-hairpin peptides derived from Aβ_l7-36_ (Figure S1).^8l,82,83^ At the three corners of these triangular trimers, Leu_17_ of one monomer subunit is near Ala_21_ of an adjacent monomer subunit. To stabilize 2AM-L into a triangular trimer, we mutated Leu_17_ and Ala_21_ to cysteine to create 2AM-L_CC_ (Figures 1A and D). Oxidation of 2AM-L_CC_ in aqueous DMSO with triethylamine (TEA) generates 2AT-L. LC-MS analysis of the oxidation reaction mixture shows that 2AM-L_CC_ crosslinks to form two major products-2AT-L and 2AM-L_CC_ with an intramolecular disulfide bond (Figure 1F). 2AT-L is isolated from the crude reaction mixture using reverse-phase HPLC. Oxidation of −30 mg of 2AM-L_CC_ typically yields −8-10 mg 2AT-L of >98% purity.

### X-Ray Crystallographic Structure of 2AT-L

We determined the X-ray crystallographic structure of 2AT-L at 1.8-A resolution (PDB 7U4P). The X-ray crystallographic structure reveals that 2AT-L is composed of three folded β-hairpins that are crosslinked together in the envisioned manner, in which Cys_17_ on one monomer forms a disulfide bond with Cys_21_ of the adjacent monomer at each corner (Figure 2A). The Aβ_17-23_ and Aβ_30-36_ β-strands of the three β-hairpins that comprise 2AT-L consist mainly of residues from the hydrophobic central and *C*-terminal regions of Aβ, creating two hydrophobic surfaces on 2AT-L (Figure 2B). The three hydrophilic Aβ_24-29_ loops extend off the hydrophobic core of 2AT-L.

**Figure 2.**
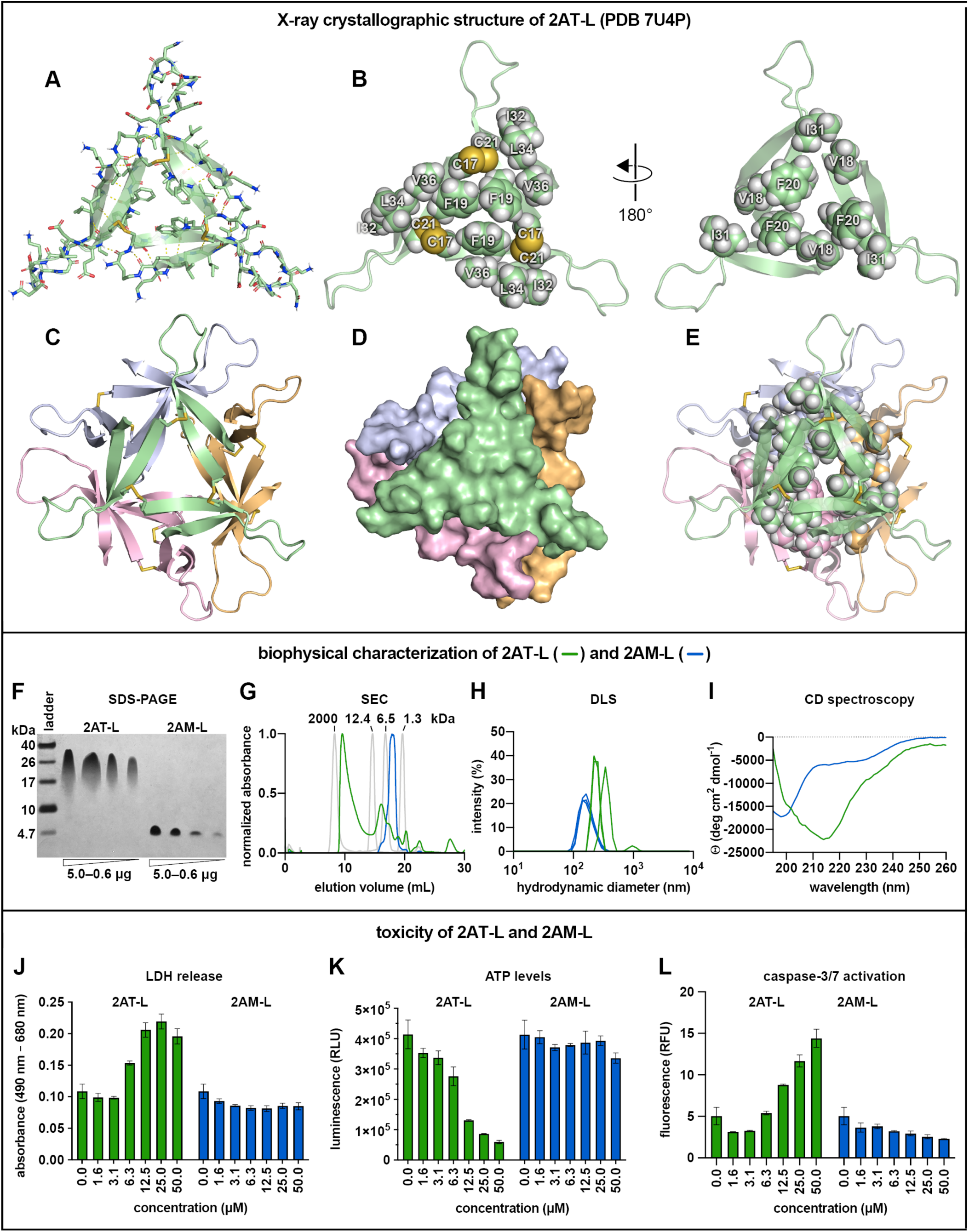
Structural, biophysical, and cell-based toxicity studies of 2AT-L. ***(A)*** X-ray crystallographic structure of 2AT-L illustrating the three folded Aβ_l7-36_ β-hairpins that comprise 2AT-L (PDB 7U4P). ***(B)*** Cartoon and sphere models of 2AT-L illustrating the two hydrophobic surfaces of 2AT-L and the hydrophilic loops that extend off the core of the trimer. ***(C)*** X-ray crystallographic structure of the ball-shaped dodecamer formed by four copies of 2AT-L. ***(D)*** Surface rendering of the ball-shaped dodecamer formed by 2AT-L illustrating how the four trimers fit together to form the dodecamer. ***(E)*** Cartoon and sphere model of the ball-shaped dodecamer formed by 2AT-L illustrating the hydrophobic core formed by Val_18_, Phe_20_, and Ile_31_ at the center of the dodecamer. ***(F)*** Silver stained SDS-PAGE of varying amounts of 2AT-L and 2AM-L. SDS-PAGE was performed in Tris buffer at pH 6.8 with 2% (w/v) SDS. ***(G)*** SEC chromatograms of 2AT-L and 2AM-L. SEC was performed on 1.0-mg/mL solutions of 2AT-L and 2AM-L in 50 mM Tris buffer (pH 8.0) with 150 mM NaCl using a Superdex 75 10/300 column. Dextran blue (2000 kDa), cytochrome C (12.4 kDa), aprotinin (6.5 kDa), and vitamin B_12_ (1.3 kDa) were run as size standards. ***(H)*** DLS traces of 2AT-L and 2AM-L. DLS traces were acquired on a 25 µM solution of 2AT-L and a 75 µM solution of 2AM-L in 10 mM phosphate buffer at pH 7.4 after centrifugation at 16,000 x g for 5 min. ***(l)*** CD spectra of 2AT-L and 2AM-L. CD spectra were acquired on a 25-µM solution of 2AT-L and a 75-µM solution of 2AM-L in 10 mM phosphate buffer at pH 7.4 ***(J)*** LDH release assay of 2AT-L and 2AM-L. ***(K)*** CellTiter-Glo ATP assay of 2AT-L and 2AM-L. ***(L)*** Caspase-3/7 activation assay of 2AT-L and 2AM-L. The assays in J-L were performed by exposing SH-SY5Y cells (30,000 cells/well on a black-walled half-area 96-well plate) to a twofold dilution series of 2AT-L and 2AM-L (50 µM to 1.6 µM) for 72 h. Each assay was performed according to manufacturer’s instructions. Data from these assays are shown as the mean of three technical replicates, with error bars representing the standard deviation.

In the crystal lattice, four copies of 2AT-L assemble to form a ball-shaped dodecamer (Figures 2C and D). The dodecamer is stabilized by an edge-to-edge hydrogen-bonding network between the backbones of adjacent trimers, and by hydrophobic packing at the core of the dodecamer between the surfaces of the trimers that contain Val_18_, Phe_20_, and Ile_31_ (Figure 2E). In total, the dodecamer contains 34 intermolecular hydrogen bonds between the four copies of 2AT-L, and the core is packed with 36 hydrophobic amino acid side chains. The outer surface of the dodecamer displays the hydrophobic amino acids on the other surface of the trimer-Phe_19_, Ile_32_, Leu_34_, and Val_36_, as well as the disulfide bond between Cys_17_ and Cys_21_. The propensity to form dodecamers appears to be a common characteristic of triangular trimers derived from Aβ_17-36_, as we have observed similar dodecameric assemblies in previous studies.^79,80,81,82,83^

### Biophysical Studies of 2AT-L

To investigate the structure and assembly of 2AT-L in solution, we turned to SDS-PAGE, size exclusion chromatography (SEC), dynamic light scattering (DLS), and circular dichroism (CD) spectroscopy. SDS-PAGE reveals that in the membrane-like environment of SDS micelles, 2AT-L assembles to form a dodecamer (Figure 2F). In SDS-PAGE, 2AT-L migrates just above the 26-kDa molecular weight marker, which is consistent with the molecular weight of a dodecamer (−26 kDa). The dodecamer band of 2AT-L is comet-shaped and streaks downward, indicating that under the conditions of SDS-PAGE the dodecamer is in equilibrium with smaller assemblies, such as hexamers and nonamers. The streaks become fainter at lower concentrations, suggesting that formation of lower-order oligomers by 2AT-L is concentration dependent; however, this could also reflect the sensitivity of the silver stain. 2AM-L migrates at or below the 4.7-kDa molecular weight marker, which is consistent with the molecular weight of a monomer or dimer. The assembly of 2AT-L to form a dodecamer in SDS-PAGE is consistent with the observation of the ball-shaped dodecamer in the crystal lattice of 2AT-L, suggesting that the ball-shaped dodecamer is the actual assembly that 2AT-L forms in a membrane-like environment and is not merely an artifact of crystal lattice formation.

To investigate the assembly of 2AT-L in an aqueous environment in the absence of SDS, we used SEC and DLS. For SEC, we ran 2AT-L on a Superdex 75 column and eluted with TBS (50 mM Tris buffer at pH 8.0 with 150 mM NaCl). Under these conditions, 2AT-L primarily elutes at 9.6 mL, indicating that 2AT-L assembles to form large species, ca. 10^2^-10^3^ kDa, well above the 26-kDa size of a dodecamer (Figure 2G). The elution profile for 2AT-L also shows a minor peak at 16.3 mL between the 12.4-kDa and 6.5-kDa size standards, which is consistent with the molecular weight of the trimer itself. Investigation of 2AT-L using DLS, shows that in phosphate buffer (10 mM sodium phosphate at pH 7.4) 2AT-L forms large species with hydrodynamic diameters of ca. 300 nm (Figure 2H). The SDS-PAGE, SEC, and DLS experiments support an assembly model where in aqueous solution, 2AT-L aggregates to form large species, and SDS dissociates these large species into their component parts, which appear to be dodecamers.

In SEC, 2AM-L elutes between the 1.3-kDa and 6.5-kDa size standards, which is consistent with the molecular weight of a monomer or dimer (Figure 2G). In contrast, in DLS, 2AM-L forms large species with hydrodynamic diameters of ca. 150 nm (Figure 2H). The different assembly properties of 2AM-L in SEC and DLS might be explained by differences in these techniques-SEC is performed under flowing conditions through a gel matrix, which may cause sheering, whereas DLS is performed in a still solution with no matrix.

To better understand the structures of 2AT-L and the higher-order assemblies formed by 2AT-L in solution, we used CD spectroscopy. In phosphate buffer, the CD spectrum of 2AT-L shows a minimum centered at 218 nm, which is characteristic of β-hairpins (Figure 2I).^84,85,86^ In contrast, the CD spectrum of 2AM-L shows a minimum near 200 nm, with shallow negative ellipticity from ca. 210-240 nm, which suggests random coil structure. These data support a structural model in which 2AT-L and the higher-order assemblies formed by 2AT-L are composed of folded β-hairpins, but in which 2AM-L does not fold to form a β-hairpin. These contrasting behaviors of 2AT-L and 2AM-L demonstrate the cooperativity between folding and assembly often observed for amyloidogenic peptides and proteins.^78,79^ Furthermore, the CD data suggest that in solution, the component β-hairpin peptides of 2AT-L adopt the folded conformation observed in the X-ray crystallographic structure of 2AT-L.

### Cell-Based Toxicity Studies of 2AT-L

Oligomers of full-length Aβ are toxic toward cells in culture.^27,29^ To determine if 2AT-L is also toxic, we exposed the human neuroblastoma cell line SH-SY5Y to 2AT-L and assessed three different metrics of toxicity: LDH release, ATP reduction, and caspase-3/7 activation. In each of the three assays, we first exposed SH-SY5Y cells to varying concentrations of 2AT-L or 2AM-L (0-50 µM) for 72 hours before performing the assay. The three toxicity metrics indicate that 2AT-L is toxic toward SH-SY5Y cells in a dose-dependent manner (Figures 2J-L). Exposing the SH-SY5Y cells to 2AT-L increased LDH release and reduced ATP levels at concentrations as low as 6.3 µM, and activated caspase-3/7 at concentrations as low as 12.5 µM. In contrast, exposing SH-SY5Y cells to the monomer 2AM-L caused little to no change in any of the three toxicity markers at concentrations up to 50 µM, which is equivalent to 16.7 µM of the trimer 2AT-L.

The findings from the structural, biophysical, and toxicity studies of 2AT-L indicate that 2AT-L behaves like an Aβ oligomer. 2AT-L assembles in the crystal lattice and in the membrane-like environment of SDS micelles to form a dodecamer. SDS-stable Aβ dodecamers, composed of antiparallel β-sheets have been observed in protein extracts from mouse and human brains.^33,34,121^ The dodecamer formed by 2AT-L may serve as a structural model for these Aβ dodecamers. The large assemblies formed by 2AT-L in the aqueous environments of SEC and DLS recapitulate previously observed large assemblies of full-length Aβ.^31,87,88^ Furthermore, like oligomers of full-length Aβ, 2AT-L is toxic toward cells in culture. The toxicity studies suggest that 2AT-L elicits toxicity by interacting with the cells and promoting membrane disruption and release of LDH, depleting ATP, and activating caspase-3/7-mediated apoptosis. Recently, the identification, characterization, and study of the putative Aβ dodecamer Aβ*56 has been called into question.^89,90^ While we do not know whether Aβ*56 is real or an artifact, the formation of dodecamers in the crystal state and in SDS-PAGE by 2AT-L and other Aβ β-hairpin peptides demonstrates that peptides derived from Aβ have a propensity to form dodecamers and that dodecamers are inherently stable structures.

### Generation and selectivity of antibodies against 2AT-L

While the structural, biophysical, and cell-based studies described above show that 2AT-L behaves like an Aβ oligomer, these studies do not on their own establish a relationship between 2AT-L and biogenic assemblies of full-length Aβ formed in the brain. To investigate the relationship between 2AT-L and Aβ assemblies that form in the brain, we generated a polyclonal antibody (pAb) against 2AT-L (pAb_2AT-L_) and then examined the immunoreactivity of this antibody with post-mortem brain tissue from people who lived with Alzheimer’s disease and people who lived with Down syndrome, as well as brain tissue from 5xFAD transgenic mice. The goal of these studies was to determine if antibodies raised against the synthetic Aβ oligomer model 2AT-L recognize biogenic Aβ assemblies, and thus provide evidence that 2AT-L may share structural or conformational epitopes with assemblies of full-length Aβ.

To generate pAb_2AT-L_, 2AT-L was first conjugated to the carrier protein keyhole limpet hemocyanin (KLH), and then rabbits were immunized with the trimer-KLH conjugate in Freunds adjuvant. Antibody titers in the rabbits reached high levels after two immunizations and remained high with repeated boosts over the course of the immunization schedule. We purified pAb_2AT-L_ from rabbit blood plasma by affinity chromatography using 2AT-L conjugated to NHS-activated agarose. The affinity-purified pAb_2AT-L_ was used in all subsequent studies.

The Aβ oligomer model 2AT-L has unique conformations, multivalency, and structures that are not present on the monomer 2AM-L; conversely, 2AT-L shares significant sequence homology with 2AM-L. Thus, 2AT-L displays unique epitopes that are not present on 2AM-L, as well as epitopes that are not unique and are present on 2AM-L. To investigate the selectivity of pAb_2AT-L_ for epitopes that are unique to 2AT-L, we compared the binding of pAb_2AT-L_ to 2AT-L and the corresponding monomer 2AM-L using an indirect ELISA. In this ELISA experiment, each well of a 96-well plate was treated with 50 ng of either 2AT-L or 2AM-L, or 1% bovine serum albumin (BSA) as a negative control. A three-fold dilution series of pAb_2AT-L_ was then applied to the wells, followed by an HRP-conjugated anti-rabbit IgG secondary antibody. The ELISA showed that pAb_2AT-L_ binds 2AT-L with a half-maximal effective concentration (EC_50_) of 0.02 μg/mL, while it only binds 2AM-L with an EC_50_ of 0.13 μg/mL (Figure S2). Thus, pAb_2AT-L_ is 6.5-fold more selective for 2AT-L than for 2AM-L. The greater selectivity for 2AT-L demonstrates that pAb_2AT-L_ is more selective for epitopes unique to the triangular trimer 2AT-L than epitopes shared by 2AT-L and the monomer 2AM-L.

### lmmunoreactivity of pAb_2AT-L_ with brain tissue from people who lived with Alzheimer’s disease and people who lived with Down syndrome

Accumulation of Aβ is etiologically associated with Alzheimer’s disease and other amyloid-related diseases. ^91^ In individuals with late-onset Alzheimer’s disease (LOAD)-the most common form of the disease-Aβ oligomer levels begin to rise and plaque deposition typically starts about two decades before the onset of symptoms, and continues throughout the disease.^34,92,93,94,95^ Individuals with trisomy 21 (Down syndrome) have an additional copy of the *APP* gene, which encodes the amyloid precursor protein from which Aβ is cleaved. As a result, Aβ accumulation and subsequent plaque formation occurs much earlier in individuals with trisomy 21, with almost all having plaque pathology by 40 years of age, and many Down syndrome Alzheimer’s disease (DSAD) individuals showing clinical signs of dementia after 50 years of age.^96,97,98,99^ In individuals with cerebral amyloid angiopathy (CAA), another neuropathology often associated with Alzheimer’s disease, Aβ assemblies accumulate around arterioles and capillaries in the cerebral cortex.^100,101,102^ Although CAA and Alzheimer’s disease can occur independently, the deposition of Aβ in CAA is thought to occur concurrently with Aβ plaque deposition and contribute to dementia in Alzheimer’s disease.^103^

To explore the relationship between the trimer 2AT-L and biogenic Aβ assemblies formed in brains from people with Alzheimer’s disease, we performed immunohistochemical experiments with pAb_2AT-L_ on clinically characterized brain tissue from elderly LOAD individuals, younger DSAD individuals, and elderly LOAD individuals with CAA. Table 1 summarizes the demographics of each individual.

**Table 1.** Individual demographics.

|  |  | age | sex | PMI (hours) | NPDx1 | tangle stage | plaque stage |
| --- | --- | --- | --- | --- | --- | --- | --- |
| LOAD individuals | 1 | 90 | F | 6.58 | Alzheimer's disease | stage 6 | stage C |
|  | 2 | 90 | M | 4.17 | Alzheimer's disease | stage 6 | stage C |
|  | 3 | 89 | F | 4.08 | Alzheimer's disease | stage 6 | stage C |
| DSAD individuals | 1 | 47 | F | 6.5 | Trisomy 21 (AD present) | stage 6 | stage C |
|  | 2 | 57 | M | 4.25 | Trisomy 21 (AD present) | stage 6 | stage C |
|  | 3 | 49 | M | 6.03 | Trisomy 21 (AD present) | stage 6 | stage C |
|  | 4 | 55 | F | 4.58 | Trisomy 21 (AD present) | stage 6 | stage C |
| CAA individual | 1 | 90 | M | 3.75 | Alzheimer's disease | stage 6 | stage B |

Abbreviations: post-mortem interval (PMI), neuropathological diagnosis (NPDx1), Alzheimer's disease (AD).

To investigate the immunoreactivity of pAb_2AT-L_ with Aβ plaques from LOAD individuals, we stained brain slices from each LOAD individual with pAb_2AT-L_ and AmyTracker680, and then imaged the brain slices using confocal fluorescence microscopy. Although AmyTracker680 stained the dense cores of the plaques, no significant pAb_2AT-L_ staining was observed in or around the plaques, even after imaging at a higher laser power. (Figures 3A-C and S3 and S4). These plaques correspond to “burned-out” plaques, which are composed of only dense cores and lack the diffuse Aβ around the cores and are thought to have once been neuritic plaques.^104,105,106^

**Figure 3.**
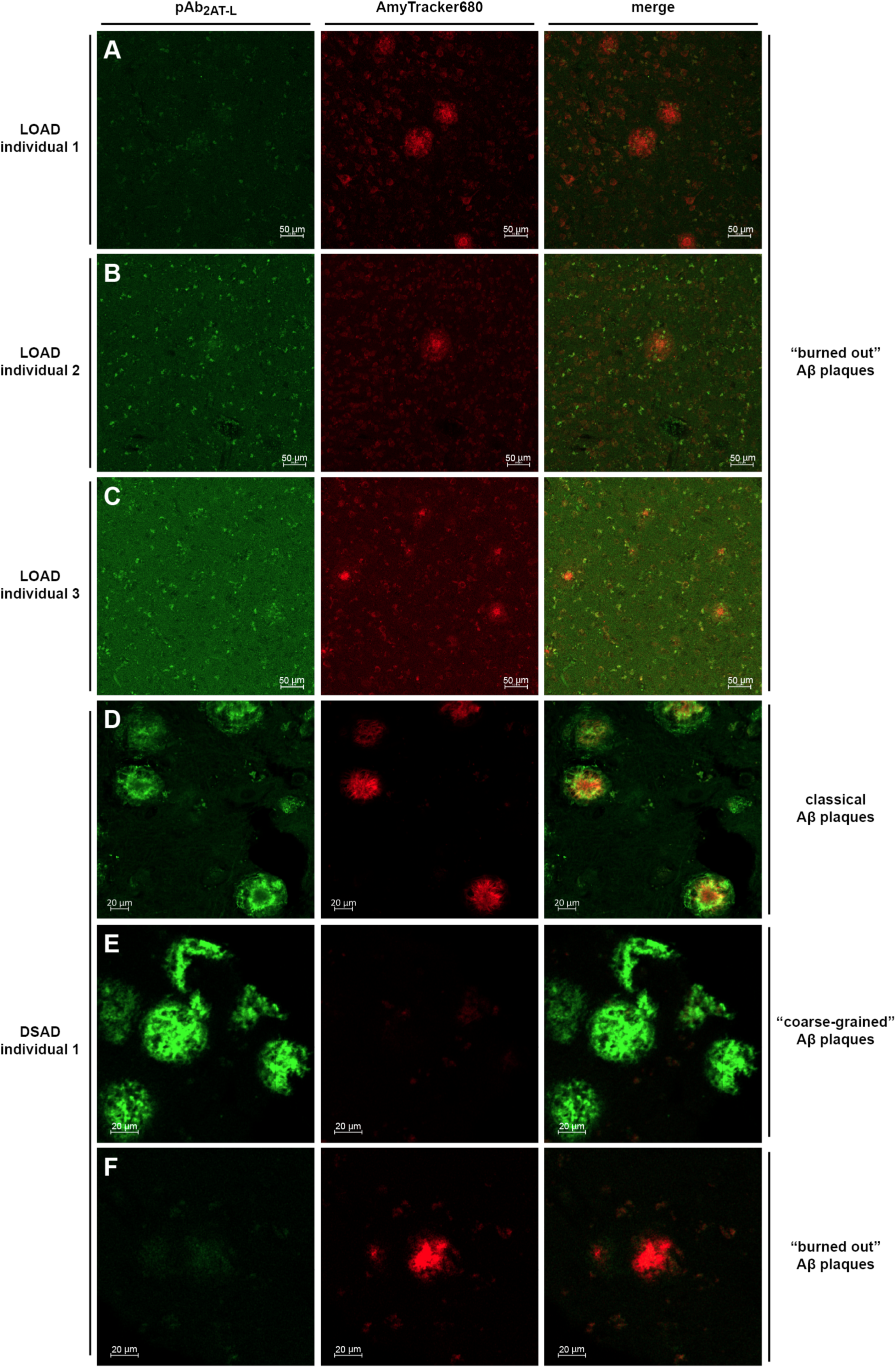
Confocal fluorescence micrographs of LOAD and DSAD brain tissue stained with pAb_2AT-L_ (green) and AmyTracker680 (red) ***(A-C)*** Representative images (10x objective) of plaques in frontal cortex brain slices from people who lived with late-onset Alzheimer’s disease (LOAD). ***(D-F)*** Representative images (20x objective) of classical Aβ plaques (D), “coarse-grained” plaques (E) and “burned-out” plaques (F) in a frontal cortex brain slice from a DSAD individual.

To investigate the immunoreactivity of pAb_2AT-L_ with Aβ plaques from the DSAD individuals, we stained a brain slice from DSAD individual 1 with pAb_2AT-L_ and AmyTracker680 and brain slices from DSAD individuals 2-4 with only pAb_2AT-L_. Confocal fluorescence microscopy reveals that pAb_2AT-L_ strongly stains plaques in the brain slices from DSAD individual 1 (Figures 3D-F) and DSAD individuals 2-4 (Figures S5-8). Three distinct plaque types that exhibit different immunohistochemical and chemical staining properties were observed in DSAD individual 1: plaques that are stained by both pAb_2AT-L_ and AmyTracker680, plaques that are only stained by AmyTracker680, and plaques that are only stained by pAb_2AT-L_.

The observation of these different plaque types is consistent with previous immunohistochemical and chemical staining studies in DSAD brain tissue slices.^107^ The plaques that are stained by both pAb_2AT-L_ and AmyTracker680 correspond to classical Aβ plaques (Figure 3D). ^108^ Classical Aβ plaques are characterized by a dense Aβ fibrillar core surrounded by more diffuse Aβ deposits that are thought to be non-fibrillar.^53,109,110,111,112,113,114^ These classical Aβ plaques show the strongest pAb_2AT-L_ staining around the peripheries of the dense cores and weaker staining of the diffuse Aβ around the dense cores, with little or no overlap of pAb_2AT-L_ and AmyTracker680 staining. The plaques that are only stained by pAb_2AT-L_ correspond to diffuse “coarse-grained” plaques, which are associated with early-onset forms of Alzheimer’s disease and are common in DSAD pathology (Figure 5E).^105,115^ The plaques that are only stained by AmyTracker680 correspond to “burned-out” dense-core plaques (Figure 5F).

The brain slices from the LOAD and DSAD individuals exhibited markedly different plaque pathologies and staining properties, an observation consistent with previous studies of LOAD and DSAD brain tissue.^107^ The LOAD tissues almost exclusively contained end-stage burned-out plaques, composed of only dense cores, which were not stained by pAb_2AT-L_. In contrast, the DSAD tissue contained multiple plaque types, many of which were strongly stained by pAb_2AT-L_. Importantly, the differences in staining between the LOAD and DSAD tissues likely does *not* reflect a preference of pAb_2AT-L_ for binding plaques in DSAD tissue over plaques in LOAD tissue, but rather, likely reflects that the DSAD tissue contains more diffuse Aβ plaques than the LOAD tissues, an observation consistent with previous studies on brain tissue from people who lived with early-onset Alzheimer’s disease and people who lived without cognitive impairment.^105,107,115,116,117,118^

To investigate the immunoreactivity of pAb_2AT-L_ with Aβ deposits in CAA, we stained brain slices from a LOAD individual exhibiting CAA pathology with pAb_2AT-L_ and AmyTracker680. Confocal fluorescence microscopy of the CAA brain slice revealed that pAb_2AT-L_ and AmyTracker680 strongly stain CAA pathology (Figure 4A). Figure 4B shows a representative image of an arteriole in which pAb_2AT-L_ and AmyTracker680 have stained Aβ deposits in the arterial walls (white arrow) and around the arteriole in the perivascular neuropil (yellow arrow). This staining and deposition pattern of Aβ is consistent with previous immunohistochemical studies of CAA brain tissue.^100–103^ The CAA tissue also contained Aβ plaques that exhibited pAb_2AT-L_ staining and AmyTracker680 staining similar to that observed in the DSAD brain slice (Figure 4C).

**Figure 4.**
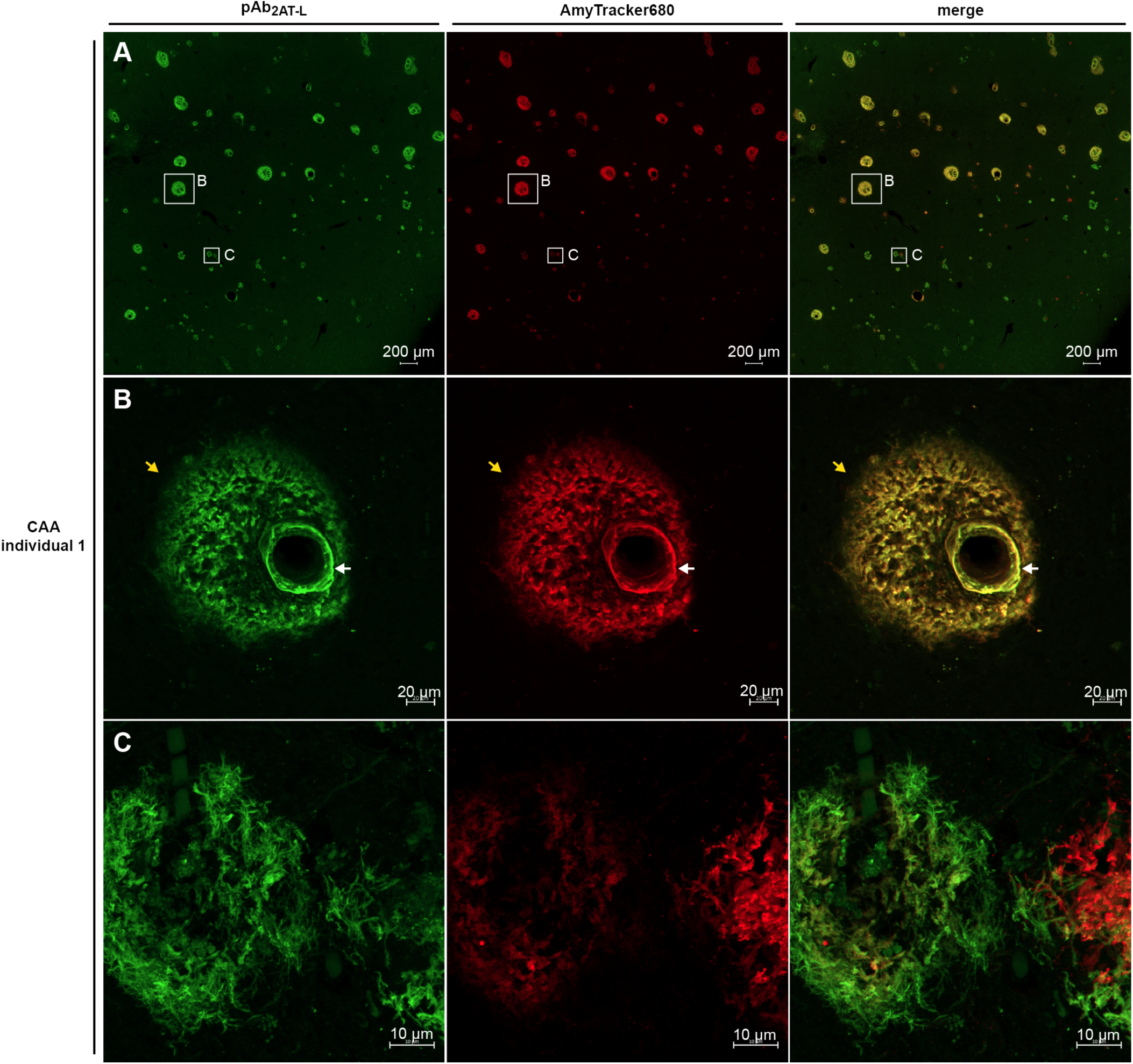
Confocal fluorescence micrographs of LOAD brain tissue containing CAA stained with pAb_2AT-L_ (green) and AmyTracker680 (red) ***(A)*** Representative stitched image (10x objective) of CAA and plaques in an occipital cortex brain slice. ***(B)*** Representative image (20x objective) of an arteriole in which pAb_2AT-L_ and AmyTracker680 have stained Aβ deposits in the arterial walls (white arrow) and around the arteriole in the perivascular neuropil (yellow arrow). ***(C)*** Representative image (63x objective) of plaques in the LOAD brain tissue containing CAA.

The CAA and Aβ plaque staining images show differences in the overlap of pAb_2AT-L_ and AmyTracker680. In the DSAD brain slices, pAb_2AT-L_ and AmyTracker680 exhibited little or no overlap in staining (Figures 3D-F). In contrast, pAb_2AT-L_ and AmyTracker680 exhibited significant overlap in staining in CAA (Figure 4). This variation is consistent with previous studies that have found that the Aβ deposits in CAA are distinct from Aβ in plaques. The 40-amino acid alloform of Aβ (Aβ_40_) predominates in CAA^119^ and is thought to form fibrils composed of parallel and anti-parallel β-sheets in CAA,^120^ while the 42-amino acid alloform of Aβ (Aβ_42_) predominates in plaques and forms fibrils composed of only parallel β-sheets. ^45,121^

The staining experiments with pAb_2AT-L_ in brain slices from individuals with Alzheimer’s disease indicate that biogenic Aβ assemblies in Alzheimer’s disease brains present epitopes that are similar to epitopes displayed on the synthetic Aβ oligomer mimic 2AT-L. These studies further support the biological significance of 2AT-L and suggest that biogenic Aβ assemblies may resemble 2AT-L. The pAb_2AT-L_ staining experiments in LOAD individuals, DSAD individuals, and LOAD individuals with CAA provide a broad overview of the immunostaining properties of pAb_2AT-L_ with Alzheimer’s disease brain tissue and indicate that antibodies raised against 2AT-L strongly recognize pathological Aβ assemblies formed in Alzheimer’s disease brains.

### lmmunoreactivity of pAb_2AT-L_ with brain tissue from 5xFAD mice

Alzheimer’s disease transgenic mouse models have aided in understanding Aβ plaque formation and its relationship to the pathogenesis and progression of Alzheimer’s disease. The Alzheimer’s disease mouse model 5xFAD contains five mutations associated with early-onset Alzheimer’s disease that lead to overproduction of Aβ_42_.^122^ 5xFAD mice exhibit accelerated Aβ plaque deposition that begins at 2 months and progresses rapidly, reaching a large plaque burden by 4-6 months and continuing to progress as the mouse ages. To explore the relationship between the trimer 2AT-L and biogenic Aβ assemblies formed in 5xFAD mouse brains, we performed immunohistochemical and immunoblotting experiments with pAb_2AT-L_ on brain tissue from 5xFAD mice.

We investigated the immunoreactivity of pAb_2AT-L_ with Aβ assemblies in 5xFAD mouse brains by staining brain slices from a 13-month-old 5xFAD mouse and a 13-month-old wild type control mouse with pAb_2AT-L_ and the amyloid-binding dye AmyTracker680 (Ebba Biotech).^123,124^ Confocal fluorescence microscopy of the 5xFAD mouse brain slice reveals that pAb_2AT-L_ binds to the outer, more diffuse Aβ deposits of the plaques (Figure 5A). Higher magnification images of representative plaques in the cortex, hippocampus, and thalamus show that the peripheries around the dense cores of the plaques exhibit the most intense staining by pAb_2AT-L_, and that the diffuse Aβ exhibits weaker, albeit still significant staining (boxed insets in Figure 5A). No staining of the dense cores by pAb_2AT-L_ was observed and no significant staining of the diffuse Aβ around the dense cores by AmyTracker680 was observed, thus there is little or no overlap in staining between pAb_2AT-L_ and AmyTracker680. No significant staining was observed in the wild type control (Figure S9).

**Figure 5.**
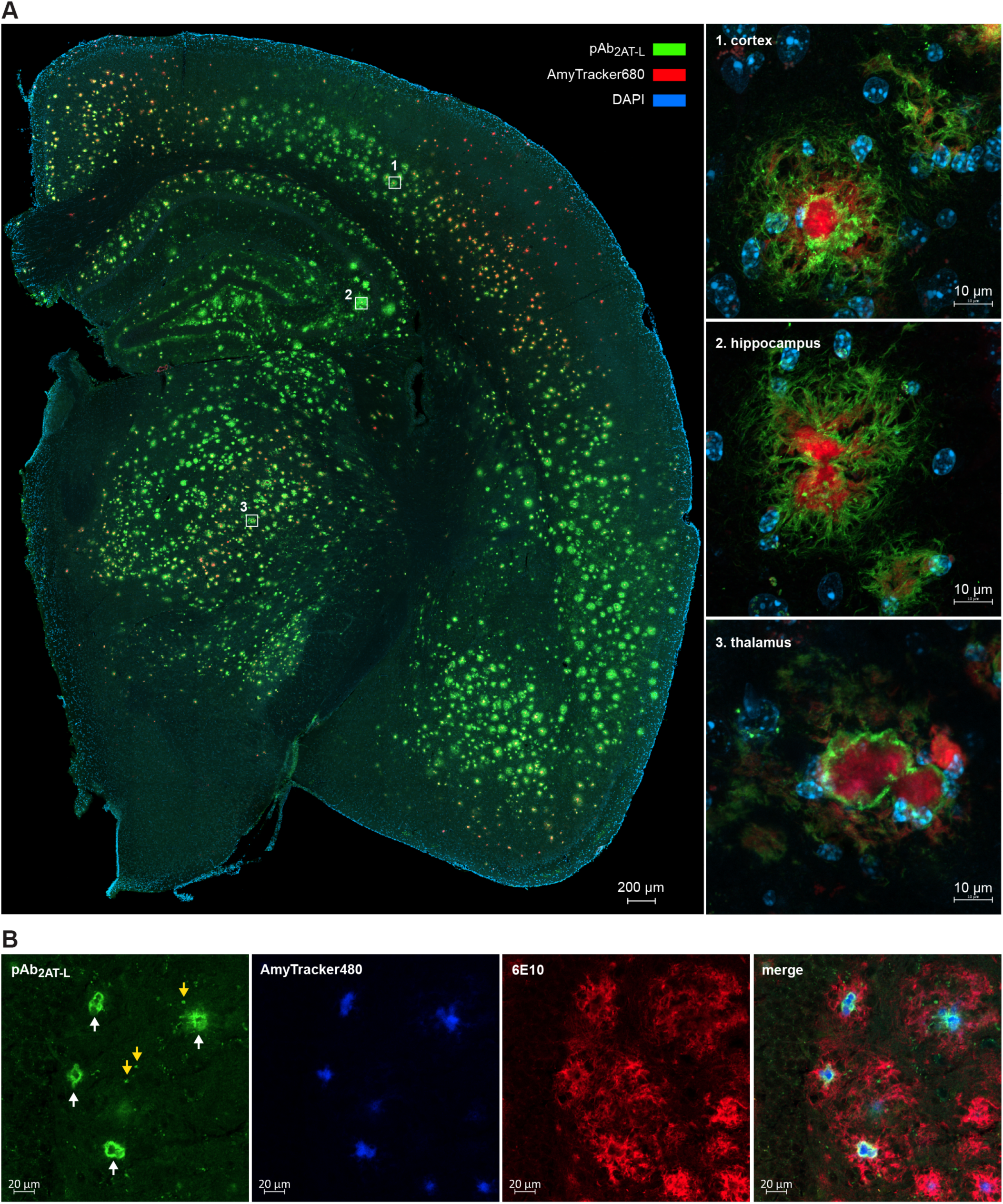
Confocal fluorescence micrographs of 5xFAD brain tissue. ***(A, left)*** Representative stitched image (1Ox objective) of a coronal brain section from a 13-month-old female 5xFAD mouse stained with pAb_2AT-L_ (green), AmyTracker68O (red), and DAPI (blue). ***(A, right)*** Representative images (63x objective) of plaques in the (1) isocortex, (2) CA3 region of hippocampus, and (3) thalamus of the 5xFAD brain slice. ***(B)*** Representative images (20x objective) of plaques in the cortex of a 13-month-old female 5xFAD mouse after extended washing after immunostaining with pAb_2AT-L_ (green) and 6E10 (red), and then subsequently staining with AmyTracker480 (blue). White arrows in the first panel designate the staining of the direct periphery of the cores by pAb_2AT-L_; yellow arrows in the first panel designate the punctate features stained by pAb_2AT-L_.

To further assess the immunostaining properties of pAb_2AT-L_ in 5xFAD mouse brain slices, we performed a subsequent experiment in which we triple labeled the plaques with pAb_2AT-L_, anti-Aβ antibody 6E10 (Biolegend), and AmyTracker480 and extended the washing step after immunostaining. In the staining experiment described in the preceding paragraph and detailed in Figure 5A, we washed the tissue 3x for 5 min in TBS with 0.1% Triton X-100 after immunostaining. In the subsequent triple-labeling experiment, we washed the tissue in TBS with 0.1% Triton X-100 overnight (−16 h) after immunostaining. Confocal fluorescence microscopy of this brain slice revealed that extended washing eliminated the weaker pAb_2AT-L_ staining of the diffuse Aβ around the dense cores, but left the pAb_2AT-L_ staining of the direct peripheries of the dense cores (white arrows in Figure 5B). The extended washing also accentuated punctate features stained by pAb_2AT-L_ that appear to reside within the diffuse Aβ around the cores (yellow arrows in Figure 5B). In contrast, the 6E10 staining of the peripheries of the dense cores and the diffuse Aβ around the dense cores is still prominent after the extended washing.

The staining experiments with pAb_2AT-L_ in 5xFAD mouse brain slices indicate that biogenic Aβ assemblies produced in 5xFAD mice present epitopes that are similar to epitopes displayed on 2AT-L, positively correlating 2AT-L with biogenic Aβ and further establishing 2AT-L as a suitable model for an Aβ oligomer. The immunostaining observed after extended washing suggests that among the antibodies in the pAb_2AT-L_ polyclonal antibody mixture, the strongest binders recognize unique features of the plaques in 5xFAD mice-the peripheries of the dense cores and punctate features embedded in the diffuse Aβ around the cores. The staining of these unique features by pAb_2AT-L_ suggests that these features are structurally distinct from the dense cores and the outer diffuse Aβ around the cores, and that pAb_2AT-L_ predominantly recognizes Aβ epitopes that are conformationally distinct from the Aβ epitopes of the diffuse Aβ around the dense cores. To our knowledge, antibodies that specifically stain the direct peripheries of the dense cores of plaques have not been previously reported.

The staining of unique features in plaques by pAb_2AT-L_ is consistent with previous findings that Aβ plaques contain structurally distinct Aβ assemblies, including Aβ oligomers.^53,125^ Ashe and coworkers isolated the dense Aβ cores and the diffuse Aβ around the cores from rTg9191 mouse brains using laser microdissection.^125^ Immunological analyses of these different plaque regions revealed that the putative Aβ dodecamer Aβ*56 and other Aβ oligomers are almost exclusively found in the diffuse Aβ of the plaques, although recent reports have called the identification, characterization, and study of Aβ*56 into question.^89,90^ Selkoe and coworkers dissolved Aβ plaques from Alzheimer’s disease individuals and used LC-MS/MS to show that the plaques contain heterogeneously cross-linked dimers of different Aβ alloforms.^30^

### lmmunoreactivity of pAb_2AT-L_ against 5xFAD brain protein extract

To corroborate that pAb_2AT-L_ recognizes biogenic Aβ in tissue, we performed biochemical experiments on brain protein extracts from 5xFAD mouse brains. We first performed a dot blot experiment to determine extraction conditions for isolating pAb_2AT-L_-reactive species. We then performed immunoprecipitation mass spectrometry experiments in which we analyzed the species pulled down by pAb_2AT-L_ by LC-MS.

To determine extraction conditions for isolating pAb_2AT-L_-reactive species we adapted a protein extraction protocol first described by Ashe and co-workers^33^ and then performed dot blot analysis on the protein extracts. In this extraction protocol, we fractionated brain proteins from a 5xFAD mouse and WT mouse into proteins soluble in TBSE (50 mM Tris buffer at pH 7.4, 100 mM NaCl, 1 mM EDTA), and proteins soluble in TBSE with detergents (TBSEd; TBSE with 3% SDS, 0.5% Triton-X, and 0.1% deoxycholate). We then spotted equal quantities of these protein extracts on nitrocellulose membranes and performed standard immunoblotting procedures with pAb_2AT-L_, 6E10, and the negative control antibody goat anti-rabbit-IgG-HRP. The dot blots show that pAb_2AT-L_ predominantly recognizes protein in the 5xFAD TBSEd fraction, showing weaker recognition of protein in the WT TBSEd extract, and little or no recognition of protein in the TBSE extracts from both 5xFAD and WT brains (Figures 6A and S10). 6E10 exhibits similar recognition properties to those of pAb_2AT-L_, showing weaker recognition toward protein in the WT TBSEd extract than pAb_2AT-L_. The goat anti-rabbit-TgG-HRP antibody exhibits no reactivity with proteins in any of the extracts.

**Figure 6.**
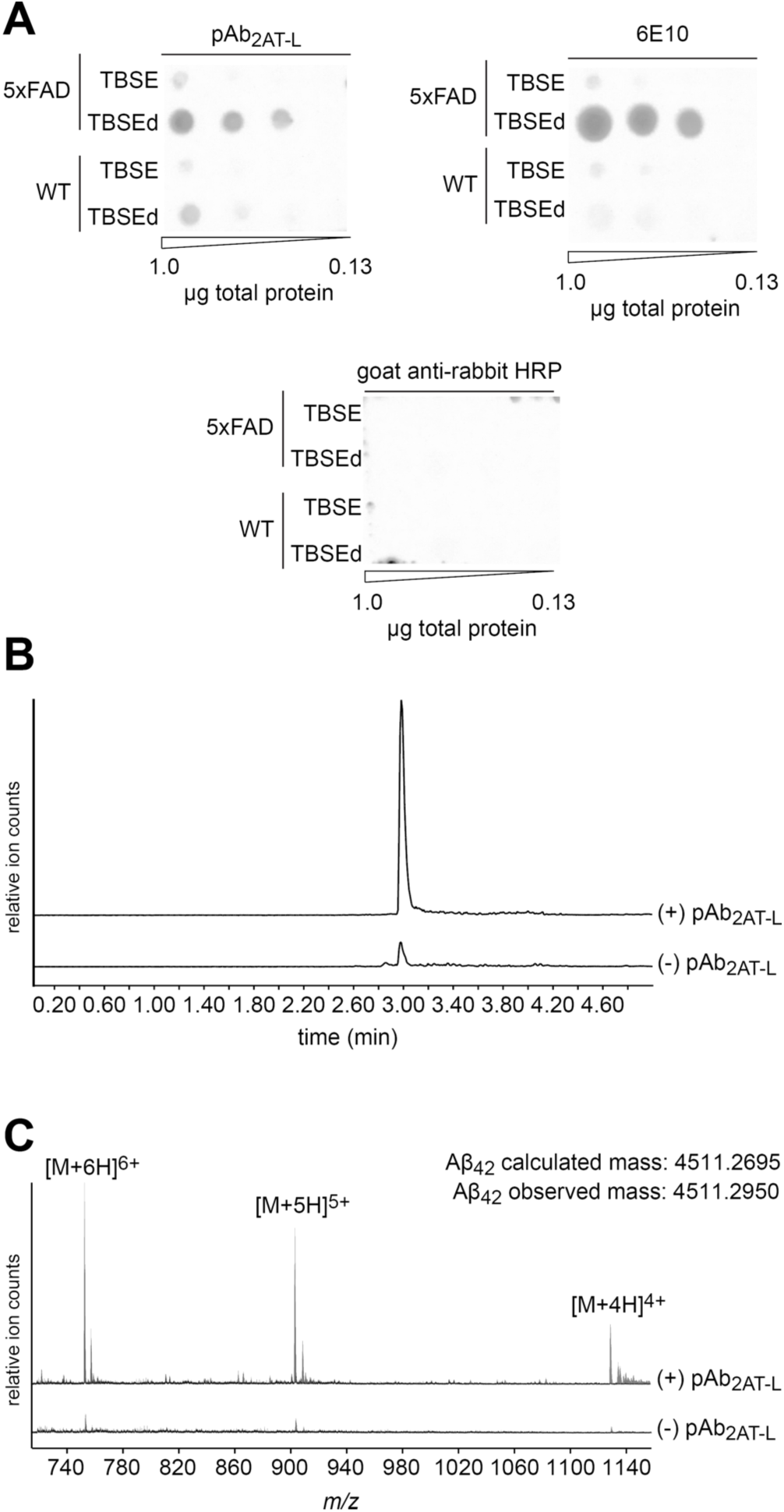
Biochemical analysis of pAb_2AT-L_ immunoreactivity with 5xFAD brain protein extracts. ***(A)*** Dot blot analysis of the immunoreactivity of pAb_2AT-L_, 6E10, and goat anti-rabbit HRP with protein extracts from 5xFAD and WT mouse brains. ***(B)*** LC-MS chromatograms of Aβ_42_ pulled down from TBSEd 5xFAD mouse brain protein extract with protein A/G Dynabeads in the presence (+) or absence (-) of pAb_2AT-L_. Chromatograms are filtered for the *m/z* of Aβ_42_. ***(CJ*** Mass spectra of the Aβ_42_ peaks from (B). The top spectrum corresponds to the top peak in (B) in which pAb_2AT-L_ was present in the pull-down experiment and shows the three Aβ_42_ charge states observed. The bottom spectrum corresponds to the bottom peak in (B) in which pAb_2AT-L_ was absent.

To determine the molecular identity of the protein species that pAb_2AT-L_ recognizes in the 5xFAD TBSEd fraction, we turned to immunoprecipitation liquid chromatography mass spectrometry (IP-LC-MS). In these experiments, we immunoprecipitated from the 5xFAD TBSEd fraction with protein A/G Dynabeads in the presence (+) or absence (-) of pAb_2AT-L_. We then washed the Dynabeads and decomplexed the bound material by treating the Dynabeads with 88% formic acid.^126,127^ Comparison of the (+) pAb_2AT-L_ and (-) pAb_2AT-L_ LC-MS chromatograms shows a peak at 2.98 min that is −10-fold more prominent in the (+) pAb_2AT-L_ sample than the (-) pAb_2AT-L_ sample (Figure 6B, Figure S10). Mass spectrometric analysis reveals that this peak is Aβ_42_ (Figure 6C, Figure S10). These results indicate that during the immunoprecipitation, pAb_2AT-L_ engages with and binds biogenic Aβ_42_ in a mixture of 5xFAD brain proteins. These results also suggest that the molecular identity of the species that pAb_2AT-L_ recognizes in the tissue staining experiments is Aβ and not another protein associated with the plaques.

## SUMMARY AND CONCLUSION

The structures of Aβ oligomers that form during Alzheimer’s disease pathogenesis and progression are unknown, constituting a significant gap in understanding the disease. Elucidating the structures of disease-relevant Aβ assemblies that form in the brain enhances our understanding of Alzheimer’s disease and holds the promise of developing better drugs that prevent or alter the course of the disease. The approach described in this paper provides a roadmap for filling this gap in understanding. This approach includes: (1) designing and synthesizing conformationally constrained Aβ β-hairpin peptides, (2) elucidating the structures of the oligomers that the Aβ β-hairpin peptides form using X-ray crystallography, (3) designing and synthesizing covalently stabilized Aβ oligomer models, (4) studying the structural, biophysical, and biological properties of the Aβ oligomer models, (5) generating antibodies against the Aβ oligomer models, and (6) characterizing the immunoreactivity of the antibodies with transgenic mouse and human brain tissue.

In this paper, we use the approach above to study the Aβ oligomer model 2AT-L, a covalently stabilized triangular trimer composed of Aβ_l7-36_ β-hairpin peptides. These studies support the biological significance of 2AT-L as an Aβ oligomer model, and suggest that Aβ assemblies that form in the brain may share structural features with 2AT-L. Structural, biophysical, and cell-based studies indicate that 2AT-L shares characteristics with oligomers formed by full-length Aβ: X-ray crystallography reveals the high-resolution structure of 2AT-L and shows that four copies of 2AT-L further assemble to form a ball-shaped dodecamer; SDS-PAGE demonstrates that 2AT-L also assembles to form a dodecamer in membrane-like environments; and cell-based studies revealed that 2AT-L is toxic toward cells. Immunohistochemical and biochemical studies with the polyclonal antibody pAb_2AT-L_ indicate that 2AT-L promotes the generation of antibodies that recognize Aβ in unique pathological features in brain tissue: immunostaining brain slices from LOAD and DSAD individuals demonstrates that pAb_2AT-L_ recognizes different types of Aβ plaques in Alzheimer’s disease brains; immunostaining of a brain slice from a LOAD individual with CAA shows that pAb_2AT-L_ recognizes Aβ that deposits around blood vessels in the brains; immunostaining of a brain slice from a 5xFAD mouse reveals that pAb_2AT-L_ recognizes Aβ around the direct peripheries of the dense cores of Aβ plaques; and immunoprecipitation LC-MS studies demonstrate that pAb_2AT-L_ engages and binds Aβ in a mixture of brain proteins and corroborates that pAb_2AT-L_ is recognizing Aβ in the immunostaining studies.

The immunoreactivity of pAb_2AT-L_ with Aβ assemblies present in plaques and CAA demonstrates that antibodies raised against 2AT-L recognize the Aβ assemblies present in these pathologies and suggests that these assemblies may share structural similarities with 2AT-L. These findings represent an important step toward understanding the structures of Aβ assemblies that form in the brain. Furthermore, these findings set the stage for pursing monoclonal antibodies against 2AT-L as well as other Aβ oligomer models our laboratory has developed.

## ASSOCIATED CONTENT

Supporting Information:

The Supporting Information is available free of charge at http://pubs.acs.org/doi/ Supporting figures, X-ray crystallography data collection and refinement statistics, materials and methods, characterization data (PDF).

Accession Codes:

Crystallographic coordinates of 2AT-L were deposited into the Protein Data Bank (PDB) with code 7U4P.

## AUTHOR INFORMATION

### Corresponding Authors

**Adam G. Kreutzer** - Department of Chemistry, University of California, Irvine, Irvine, California 92697-2025, United States; orcid.org/0000-0002-9724-6298;

**James S. Nowick** - Department of Chemistry and Department of Pharmaceutical Sciences, University of California, Irvine, Irvine, California 92697-2025, United States; orcid.org/0000-0002-2273-1029;

### Authors

**Chelsea Marie T. Parrocha** - Department of Pharmaceutical Sciences, University of California, Irvine, Irvine, California 92697-2025, United States; orcid.org/0000-0002-6502-1297

**Sepehr Haerianardakani** - Department of Chemistry, University of California, Irvine, Irvine, California 92697-2025, United States; orcid.org/0000-0003-1539-2345

**Gretchen Guaglianone** - Department of Chemistry, University of California, Irvine, Irvine, California 92697-2025, United States; orcid.org/0000-0002-5189-2550

**Jennifer T. Nguyen** - Department of Chemistry, University of California, Irvine, Irvine, California 92697-2025, United States; orcid.org/0009-0004-9748-4378

**Michelle N. Diab** - Department of Chemistry, University of California, Irvine, Irvine, California 92697-2025, United States

**William Yong** - Department of Pathology and Laboratory Medicine, University of California, Irvine, Irvine, California 92697-2025, United States

**Mari Perez-Rosendahl** - Department of Pathology and Laboratory Medicine, University of California, Irvine, Irvine, California 92697-2025, United States

**Elizabeth Head** - Department of Pathology and Laboratory Medicine, University of California, Irvine, Irvine, California 92697-2025, United States; orcid.org/0000-0003-lll5-6396

### Author Contributions

A.G.K. and J.S.N. designed the research and wrote the paper. A.G.K., C.M.T.P., S.H., G.G., J.T.N., and M.N.D. performed the peptide and trimer synthesis, purification, and characterization. A.G.K. and S.H. performed X-ray crystallography. A.G.K. performed SDS-PAGE. S.H. performed SEC, DLS, and CD spectroscopy. A.G.K. performed the cell-based toxicity studies. A.G.K. and C.M.T.P. performed the immunostaining studies. A.G.K. performed the immunoprecipitation LC-MS studies. W.Y., M. P-R., and E.H. completed neuropathology diagnoses for the Alzheimer’s disease and Down syndrome brain tissue

### Notes

The authors declare the following competing financial interest(s): The Regents of the University of California has been assigned a United States patent for compounds reported in this paper in which A.G.K. and J.S.N. are inventors.

## ACKNOWLEDGEMENTS

We thank Benjamin Katz and Dr. Felix Grun at the University of California Irvine Mass Spectrometry Facility for assistance with LC-MS, and Dr. Dmitry Fishman at the University of California Irvine Laser Spectroscopy Labs for assistance with CD spectroscopy. We thank Dr. Shimako Kawauchi, Dr. Grant MacGregor, and the staff at the University of California Irvine Transgenic Mouse Facility for breeding and genotyping the 5xFAD mice. The authors acknowledge the support of the Chao Family Comprehensive Cancer Center Transgenic Mouse Facility Shared Resource, supported by the National Cancer Institute of the National Institutes of Health under award number P30CA062203. The content is solely the responsibility of the authors and does not necessarily represent the official views of the National Institutes of Health. We thank the staff at the UCI MIND ADRC Neuropathology Core. The UCI ADRC is funded by NIH/NIA grant P30AG066519. We thank Dr. Adeela Syed at the University of California Optical Biology Core Facility for assistance with confocal microscopy. This study was made possible in part through access to the Optical Biology Core Facility of the Developmental Biology Center, a shared resource supported by the Cancer Center Support Grant (CA-62203) and Center for Complex Biological Systems Support Grant (GM-076516) at the University of California, Irvine. Beamline 5.0.1 of the Advanced Light Source, a DOE Office of Science User Facility under Contract No. DE-AC02-05CH11231, is supported in part by the ALS-ENAβLE program funded by the National Institutes of Health, National Institute of General Medical Sciences, grant P30 GM124169-01. We thank the National Institutes of Health (NIH) National Institute on Aging (NIA) for funding (Grants AG062296 and AG072587).

## FOR TABLE OF CONTENTS ONLY

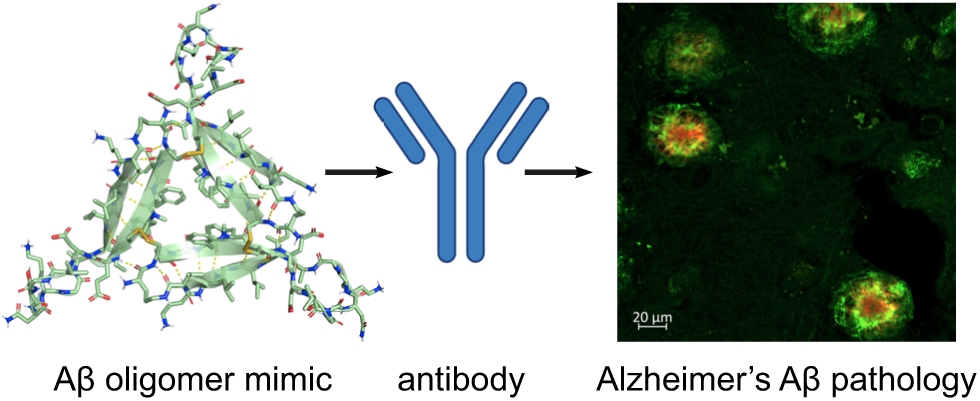

