## Supporting Information for "Antibodies Raised Against an Aβ Oligomer Mimic Recognize Pathological Features in Alzheimer’s Disease and Associated Amyloid-Disease Brain Tissue"

**This PDF includes:**

**Supporting Figures and Table**

**Figure S1.** X-ray crystallographic structures of triangular trimers composed of β-hairpin peptides derived from $A\beta_{17–36}$. S4

**Figure S2.** Indirect ELISA of $pAb_{2AT\text{-}L}$ with 2AT-L, 2AM-L and BSA. S4

**Figure S3.** Fluorescence micrograph of a brain slice from LOAD individual 1 stained with $pAb_{2AT\text{-}L}$. S5

**Figure S4.** Fluorescence micrograph of a brain slice from LOAD individual 2 stained with $pAb_{2AT\text{-}L}$. S6

**Figure S5.** Fluorescence micrograph of a brain slice from DSAD individual 1 stained with $pAb_{2AT\text{-}L}$. S7

**Figure S6.** Fluorescence micrograph of a brain slice from DSAD individual 2 stained with $pAb_{2AT\text{-}L}$. S8

**Figure S7.** Fluorescence micrograph of a brain slice from DSAD individual 3 stained with $pAb_{2AT\text{-}L}$. S9

**Figure S8.** Fluorescence micrograph of a brain slice from DSAD individual 4 stained with $pAb_{2AT\text{-}L}$. S10

**Figure S9.** Representative stitched image of a coronal brain section from a 13-month-old female wild type mouse stained with $pAb_{2AT\text{-}L}$ and DAPI. S11

**Figure S10.** IP-LC-MS data from immunoprecipitation with $pAb_{2AT\text{-}L}$ from 5xFAD mouse brain protein extract. S12

**Table S1.** Crystallographic properties, crystallization conditions, and data collection and model refinement statistics for 2AT-L. S13

**Materials and Methods**

General information. S14
Synthesis and purification of 2AM-L and 2AM-L$_{CC}$. S14
Loading of the resin. S14
Peptide coupling. S14
Cleavage of the peptide from the resin. S15
Cyclization of the linear peptide. S15
Global deprotection of the cyclic peptide. S15
Reverse-phase HPLC purification. S16
Scheme 1. Synthetic scheme for 2AM-L. S17
Synthesis, purification, and characterization of 2AT-L. S18
Synthesis of 2AT-L. S18
LC-MS analysis of the 2AM-L$_{CC}$ oxidation reaction mixture. S18
Purification of 2AT-L. S19
LC-MS characterization of 2AT-L. S20
X-ray crystallography of 2AT-L. S21
Crystallization procedure for 2AT-L. S21
X-ray crystallographic data collection, data processing, and structure determination. S21
SDS-PAGE and silver staining. S22
Sample preparation and gel running. S22
Silver staining. S22
Size exclusion chromatography (SEC). S23
Dynamic light scattering (DLS). S24
Circular dichroism (CD) spectroscopy. S24
Cell-based toxicity assays of 2AT-L and 2AM-L in SH-SY5Y cells. S24
Preparation of SH-SY5Y cells for the toxicity assays. S25
Preparation of 2AT-L and 2AM-L for toxicity assays. S25
Treatment of the SH-SY5Y cells with 2AT-L and 2AM-L. S26
CyQUANT™ LDH Cytotoxicity Assay. S26
CellTiter-Glo® 2.0 Cell Viability Assay. S26
Apo-ONE® Homogeneous Caspase-3/7 Assay. S27

Generation of $pAb_{2AT-L}$. S27
Rabbit immunization. S27
Affinity purification of $pAb_{2AT-L}$. S28
Indirect ELISA of $pAb_{2AT-L}$ against 2AT-L, 2AM-L, and BSA. S29
Coating the wells of the ELISA plate with 2AT-L, 2AM-L, and BSA. S29
Treating the ELISA plate $pAb_{2AT-L}$. S30
Treating the ELISA plate with the secondary antibody. S30
Developing the ELISA plate. S31
Immunostaining and fluorescence microscopy of human and 5xFAD mouse brain slices. S31
Preparing the human brain tissues for immunostaining. S31
Preparing the 5xFAD mouse brain tissues for immunostaining. S31
Staining and imaging the human and 5xFAD mouse brain tissue slices. S32
Preparation of 5xFAD brain protein extracts. S34
Dot blot assay of $pAb_{2AT-L}$ against 5xFAD brain protein extracts. S35
Immunoprecipitation LC-MS on 5xFAD brain protein extracts. S36
**References and Notes** S38
**Characterization Data**
Characterization of 2AM-L. S39
Characterization of 2AT-L. S41

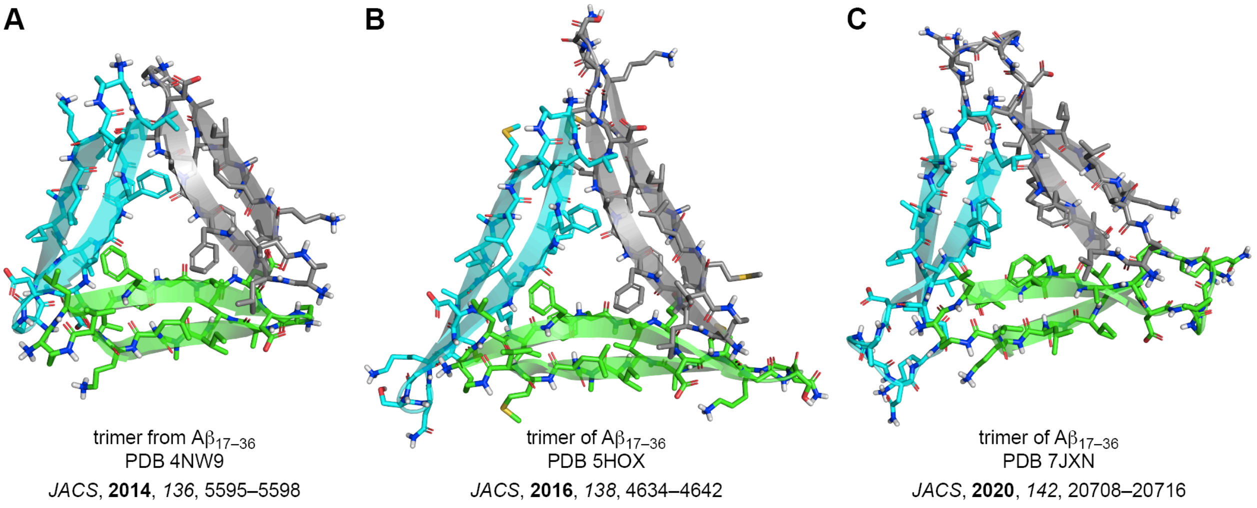


***Figure S1.*** X-ray crystallographic structures of triangular trimers composed of β-hairpin peptides derived from $A\beta_{17–36}$. ***(A)*** Trimer composed of peptides derived from an $A\beta_{17–36}$ β-hairpin that lack the $A\beta_{24–29}$ loop. ***(B)*** Trimer composed of $A\beta_{17–36}$ β-hairpins that contain an intramolecular disulfide linkage between positions 24 and 29 to reinforce β-hairpin conformation. ***(C)*** Trimer composed of $A\beta_{17–36}$ β-hairpins that contain cyclohexylalanine at position 20 in place of phenylalanine.

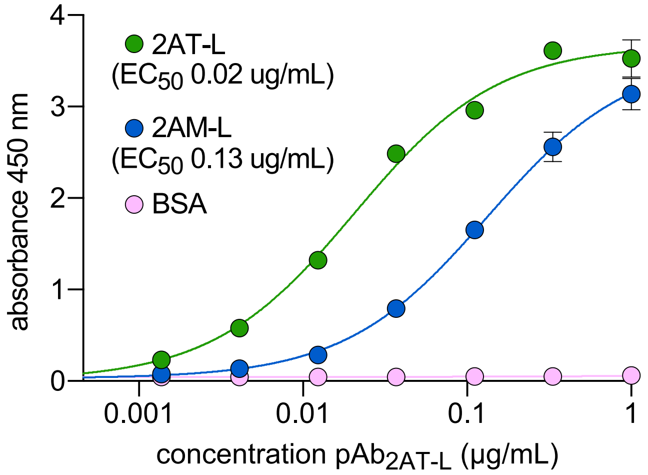


***Figure S2.*** Indirect ELISA of $pAb_{2AT-L}$ with 2AT-L, 2AM-L and BSA.

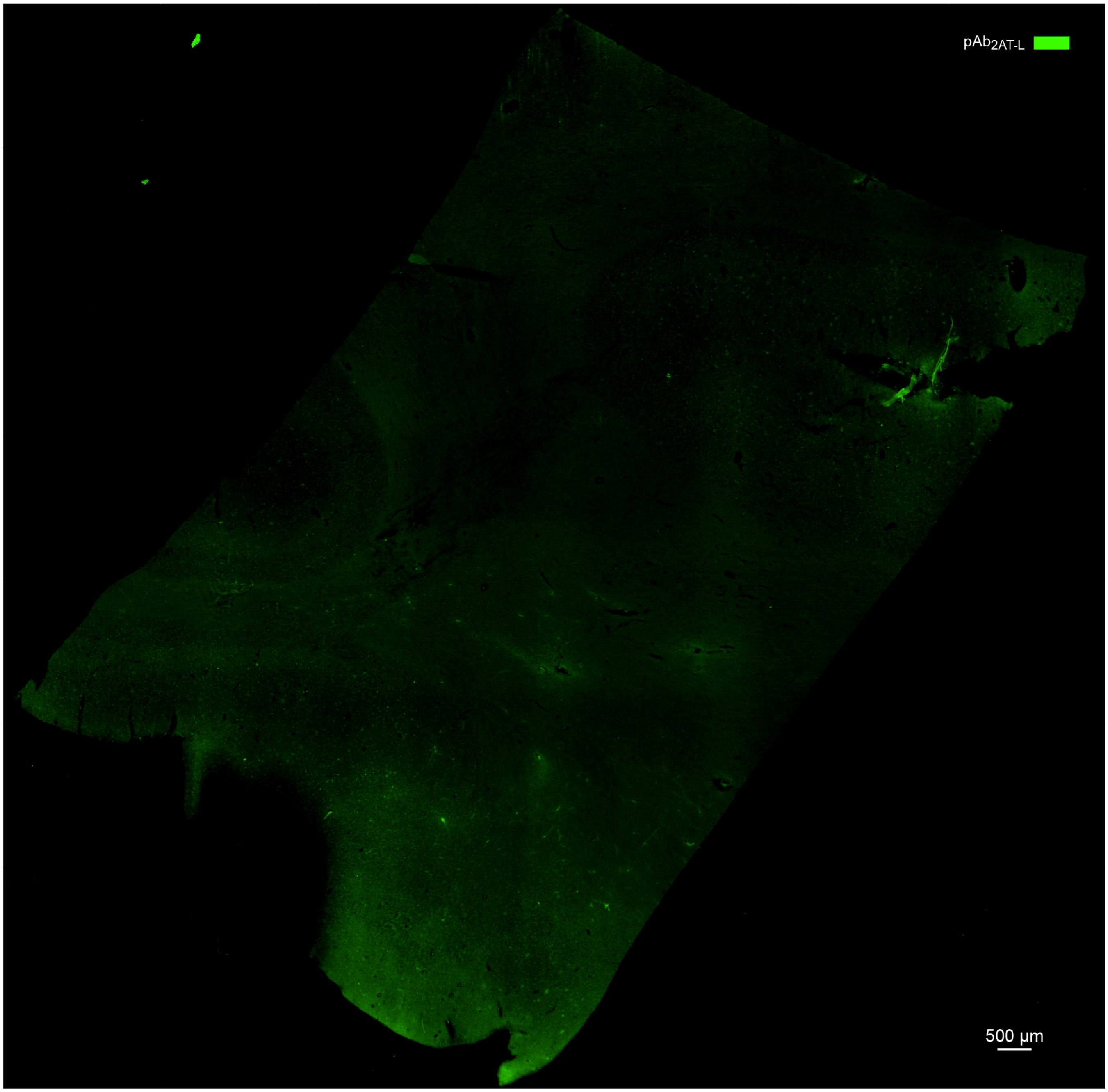


***Figure S3.*** Fluorescence micrograph (4x objective, stitched image) of a brain slice from LOAD individual 1 stained with $pAb_{2AT\text{-}L}$ (green).

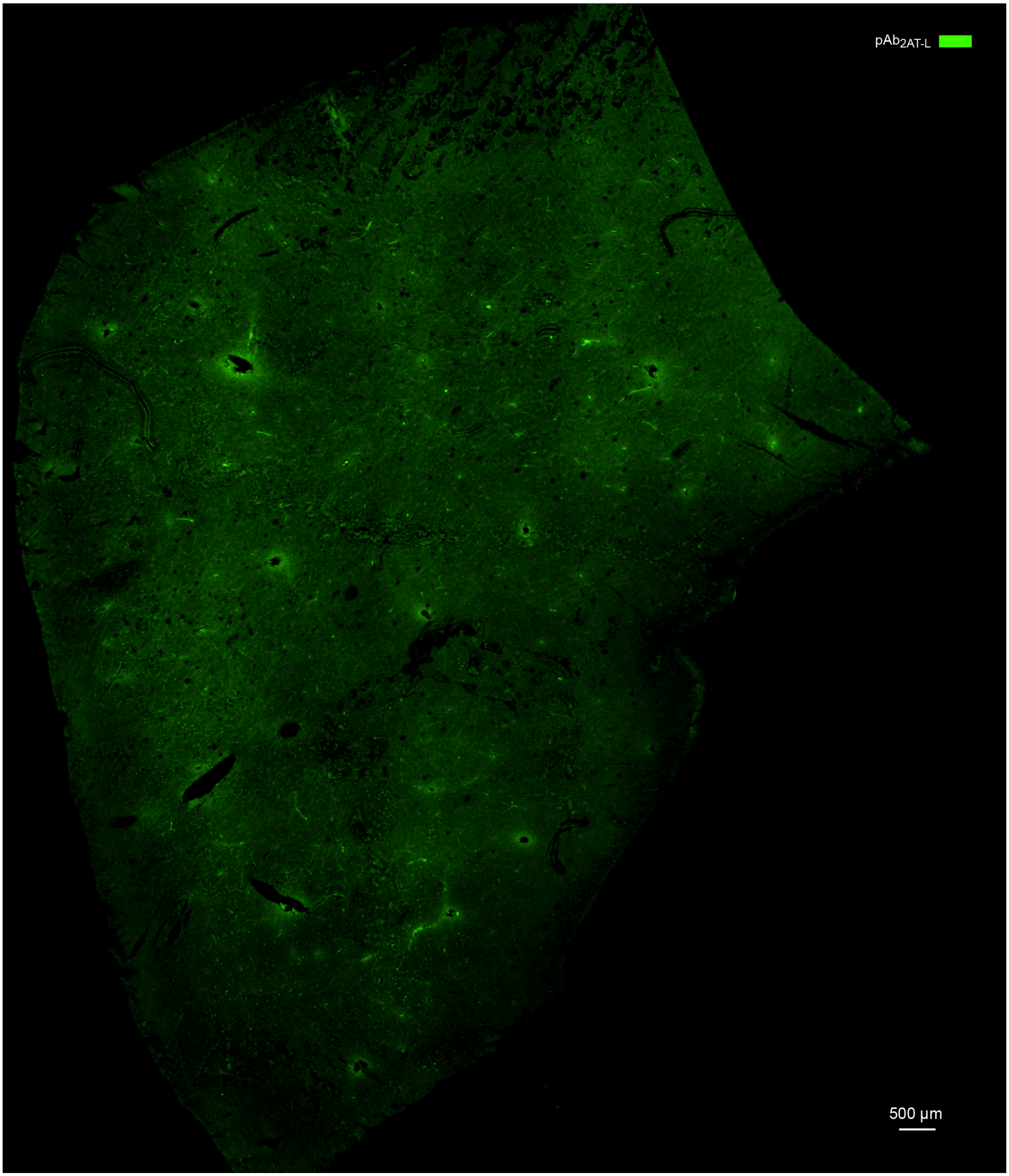


***Figure S4.*** Fluorescence micrograph (4x objective, stitched image) of a brain slice from LOAD individual 2 stained with $pAb_{2AT\text{-}L}$ (green).

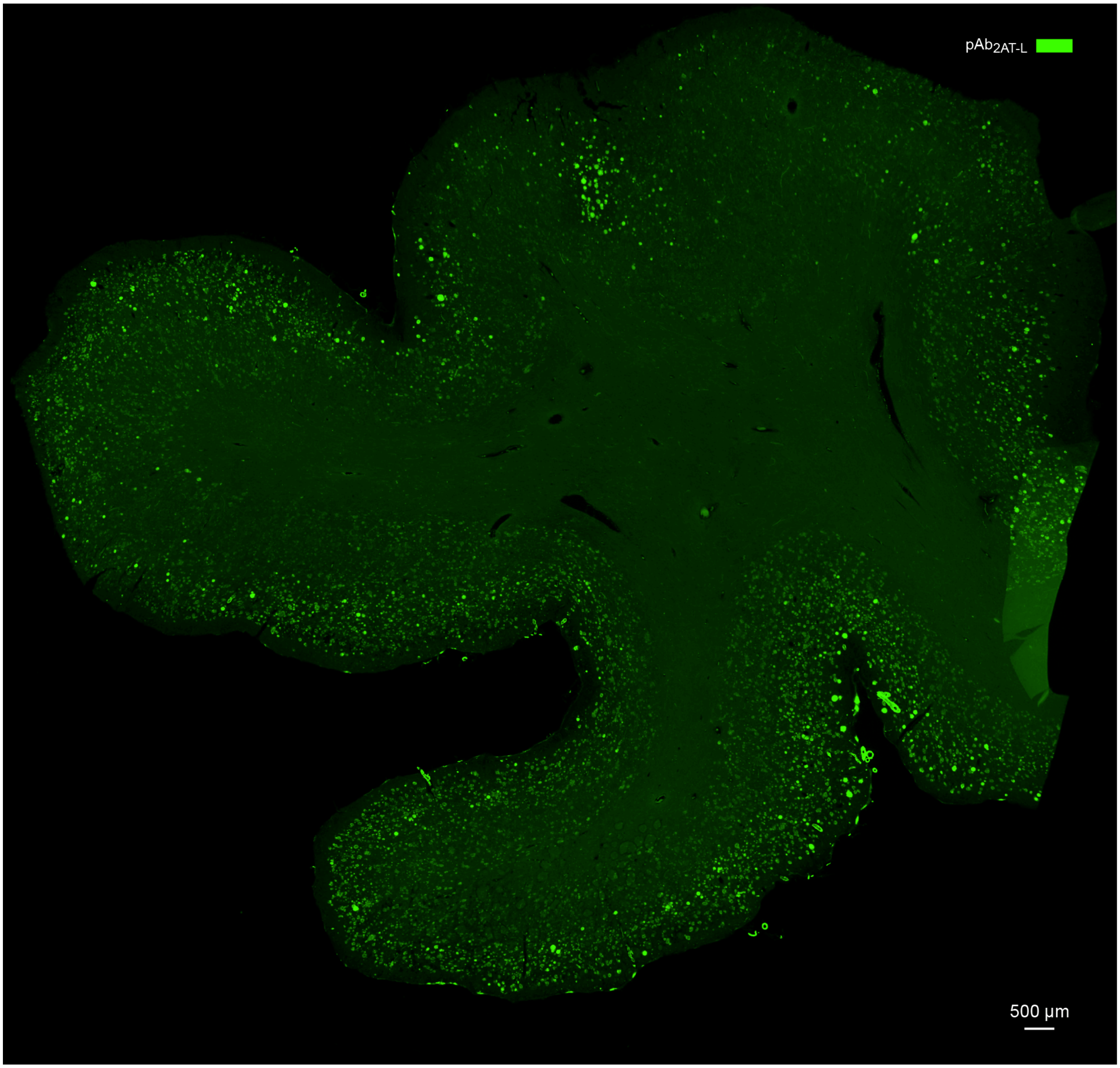


***Figure S5.*** Fluorescence micrograph (4x objective, stitched image) of a brain slice from DSAD individual 1 stained with $pAb_{2AT-L}$ (green).

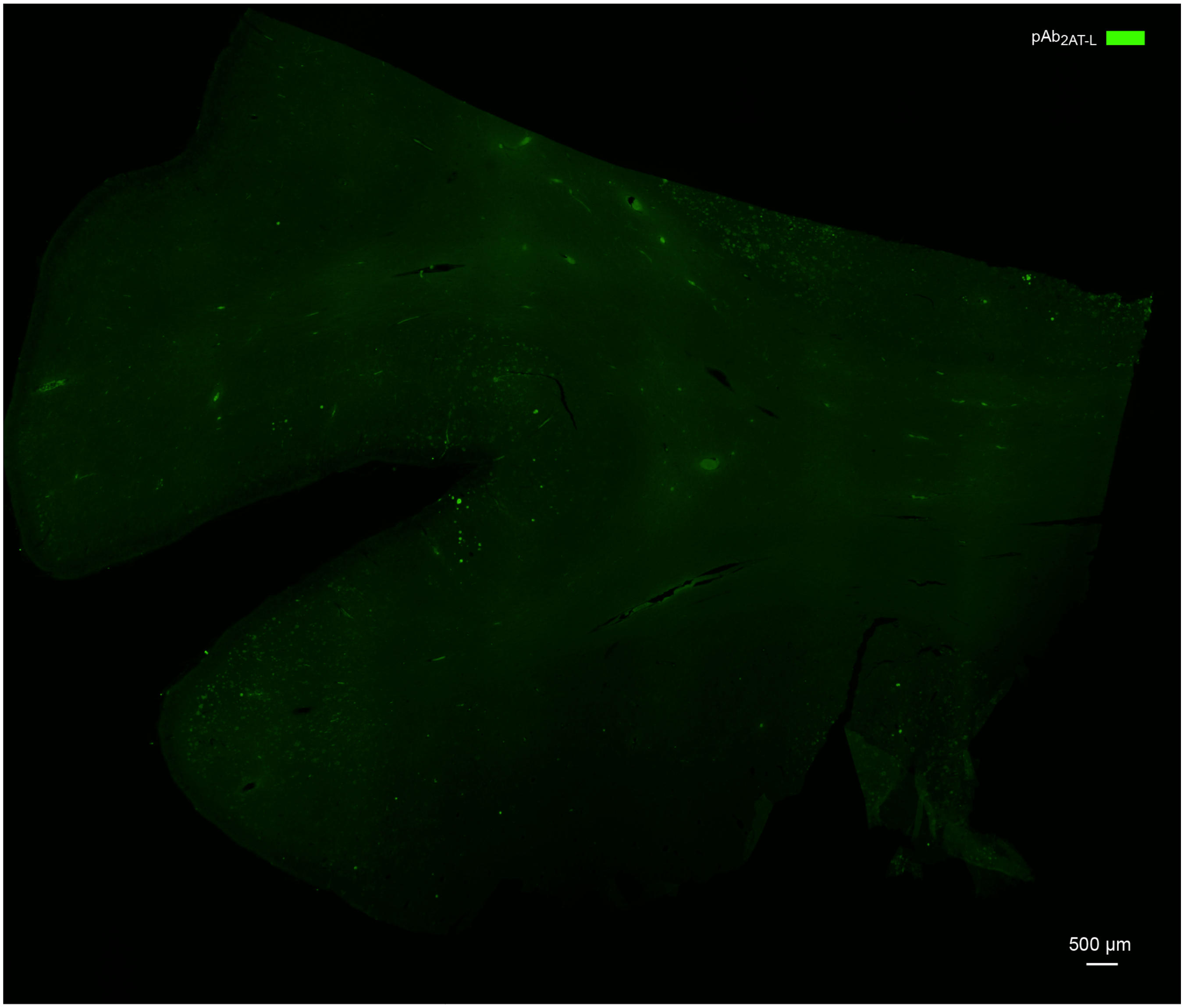


***Figure S6.*** Fluorescence micrograph (4x objective, stitched image) of a brain slice from DSAD individual 2 stained with $pAb_{2AT-L}$ (green).

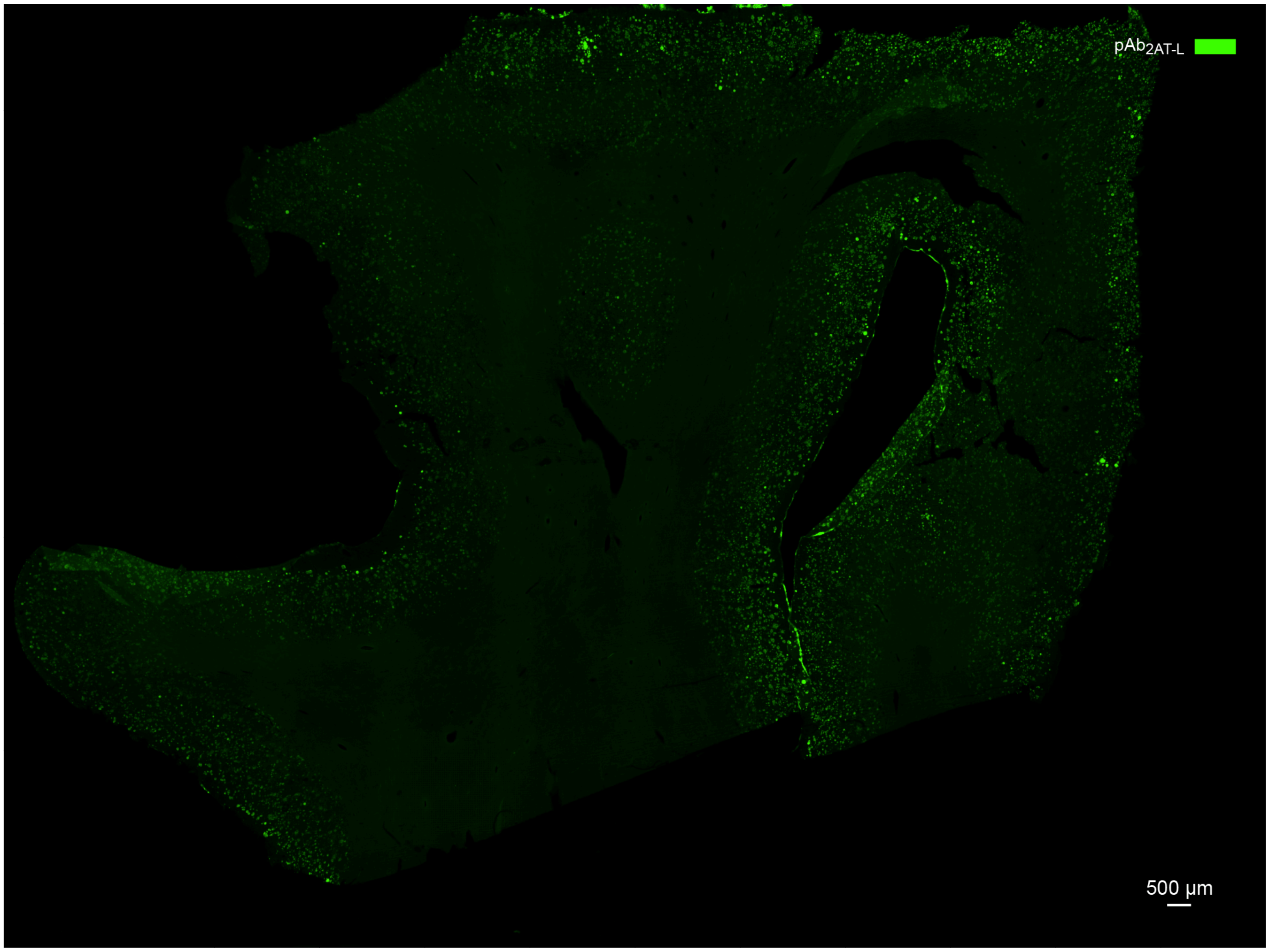


***Figure S7.*** Fluorescence micrograph (4x objective, stitched image) of a brain slice from DSAD individual 3 stained with $pAb_{2AT\text{-}L}$ (green).

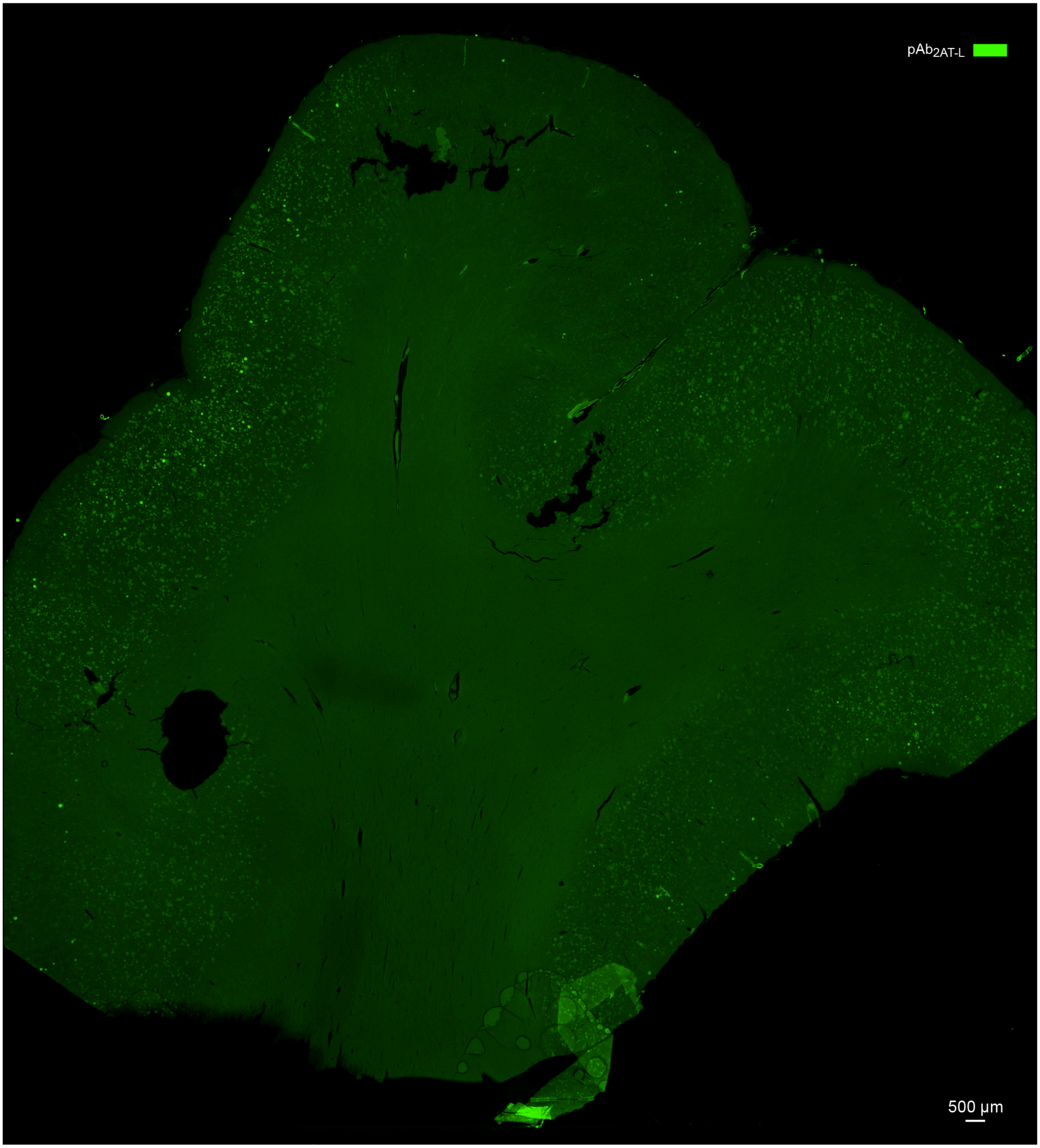


***Figure S8.*** Fluorescence micrograph (4x objective, stitched image) of a brain slice from DSAD individual 4 stained with $pAb_{2AT\text{-}L}$ (green).

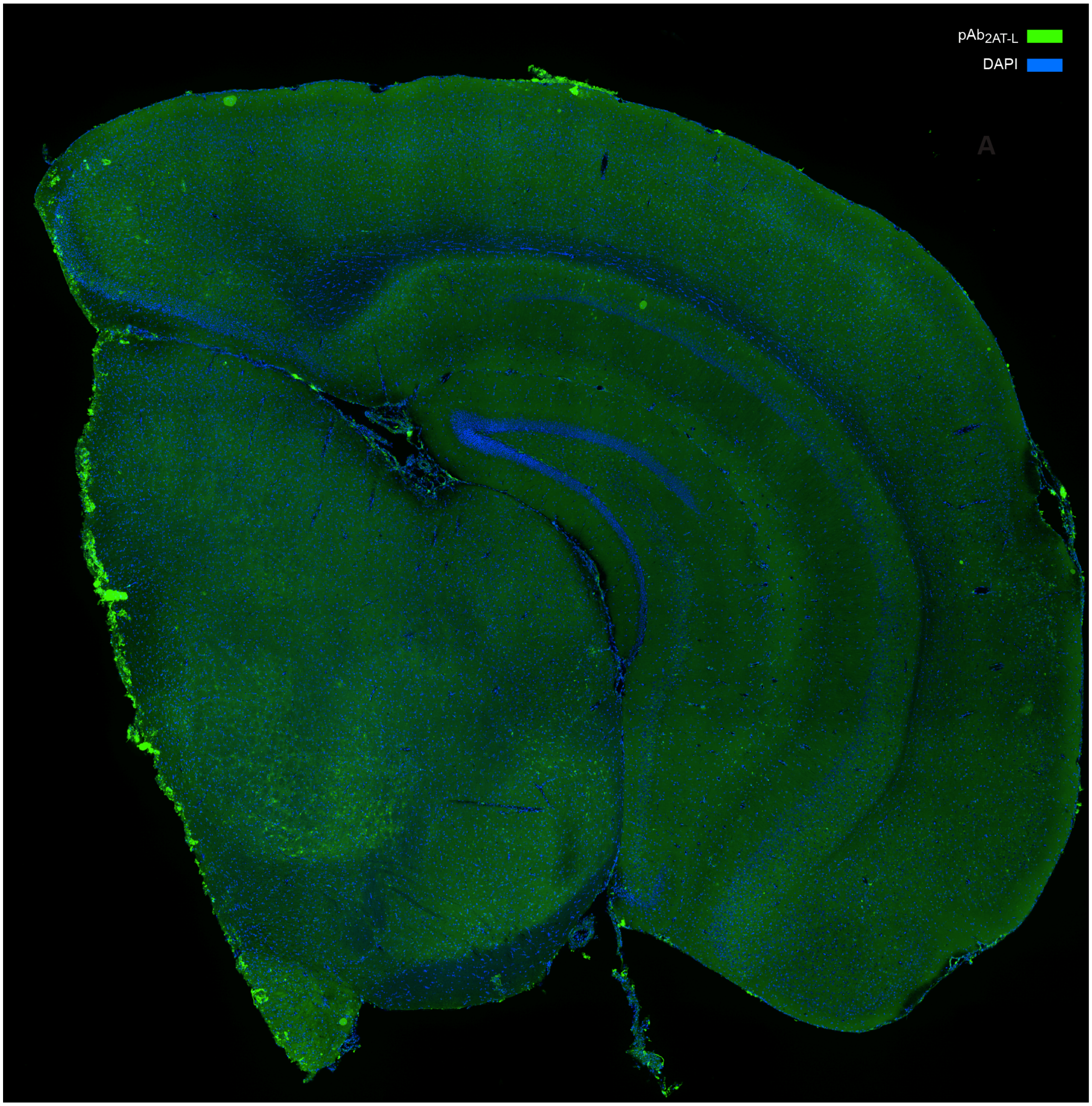


***Figure S9.*** Representative stitched image (10x objective) of a coronal brain section from a 13-month-old female wild type mouse stained with $pAb_{2AT\text{-}L}$ (green) and DAPI (blue).

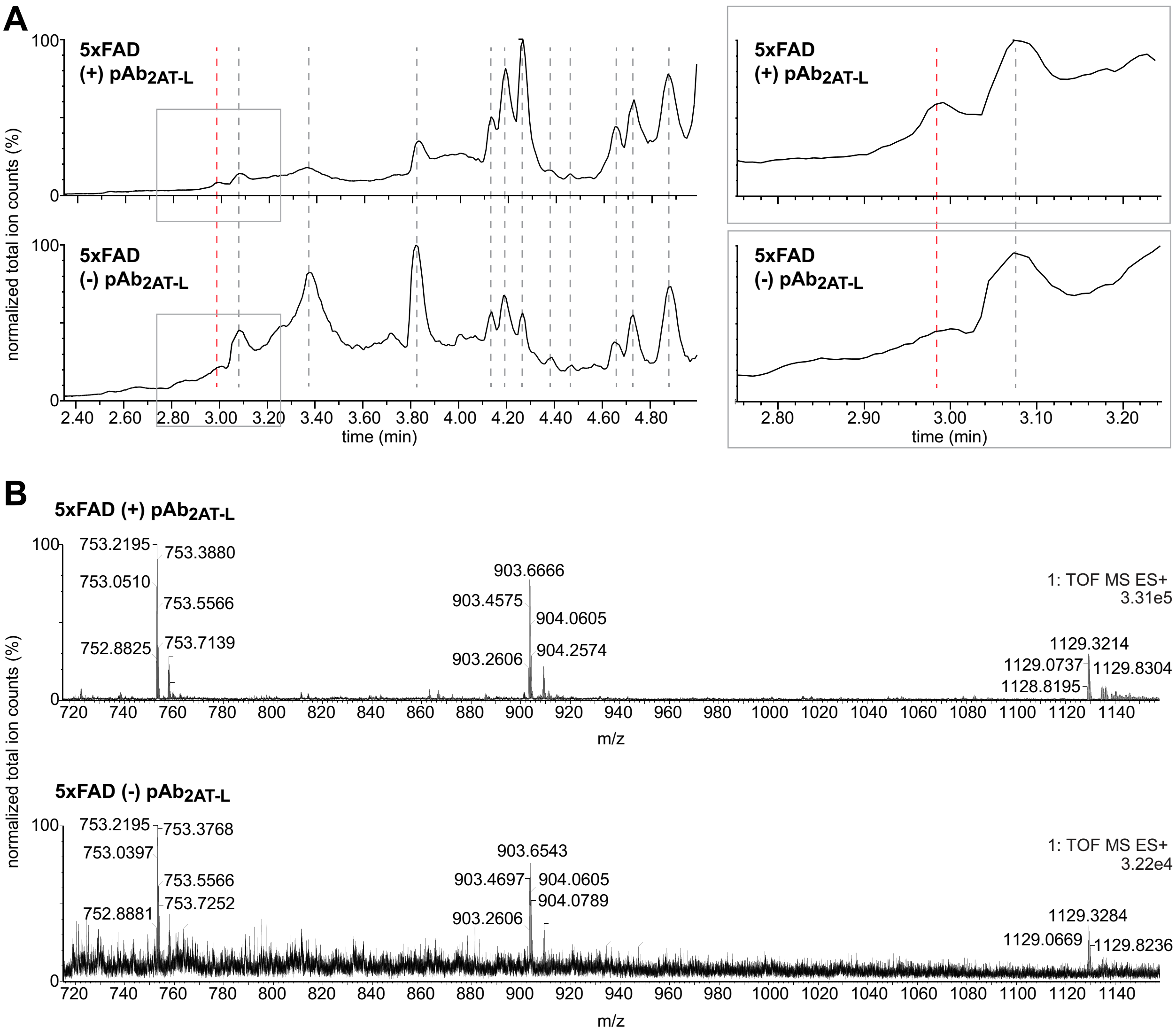


***Figure S10.*** LC-MS data from immunoprecipitation with $pAb_{2AT\text{-}L}$ from 5xFAD mouse brain protein extract. ***(A)*** LC-MS chromatograms showing all material pulled down from TBSEd 5xFAD mouse brain protein extract with protein A/G Dynabeads in the presence of $pAb_{2AT\text{-}L}$ (+) or absence of $pAb_{2AT\text{-}L}$ (-) after elution with 88% formic acid. The red dashed line designates the $A\beta_{42}$ peak. The grey boxed insets are zoomed in chromatograms showing the $A\beta_{42}$ peak ***(B)*** Mass spectra of the $A\beta_{42}$ peaks from (A).

***Table S1.*** Crystallographic properties, crystallization conditions, and data collection and model refinement statistics for 2AT-L.

| | |
|---|---|
| PDB ID | 7U4P |
| space group | $P6_222$ |
| *a*, *b*, *c* (Å) | 62.054 62.054 47.839 |
| α, β, λ (°) | 90, 90, 120 |
| molecules per asymmetric unit | 1 |
| wavelength (Å) | 0.97741 |
| resolution (Å) | 31.03–1.803 (1.868–1.803) |
| total reflections | 194477 (18796) |
| unique reflections | 5373 (522) |
| multiplicity | 36.2 (36.0) |
| completeness (%) | 99.74 (99.81) |
| mean I/σ(I) | 39.93 (4.07) |
| Wilson B factor | 31.27 |
| $R_{merge}$ | 0.0643 (0.9299) |
| $R_{measure}$ | 0.06524 (0.9431) |
| $CC_{1/2}$ | 1 (0.958) |
| $CC^*$ | 1 (0.989) |
| $R_{work}$ | 0.2054 (0.2875) |
| $R_{free}$ | 0.2475 (0.3070) |
| number of non-hydrogen atoms | 484 |
| $RMS_{bonds}$ | 0.010 |
| $RMS_{angles}$ | 1.00 |
| Ramachandran favored (%) | 92.31 |
| Ramachandran allowed (%) | 7.69 |
| Ramachandran outliers (%) | 0 |
| rotamer outliers (%) | 0 |
| clashscore | 9.23 |
| average B-factor | 56.12 |
| number of TLS groups | 6 |
| ligands/ions | 0 |
| water molecules | 114 |
| crystallization conditions | 0.1 M Tris at pH 8.3, 0.2 M $MgCl_2$, 2.8 M 1,6-hexanediol |

### Materials and Methods[1]

##### *General information*

All chemicals were used as received unless otherwise noted. Methylene chloride ($CH_2Cl_2$) was passed through alumina under nitrogen prior to use. Anhydrous, amine-free *N,N*-dimethylformamide (DMF) was purchased from Alfa Aesar. Deionized water (18 MΩ) was obtained from a Barnstead NANOpure Diamond water purification system. HPLC grade acetonitrile and deionized water, each containing 0.1% trifluoroacetic acid (TFA), were used for analytical and preparative reverse-phase HPLC. 2AM-L, 2AM-$L_{CC}$, and 2AT-L, were prepared and used as the trifluoroacetate salts and were assumed to have one trifluoroacetic acid molecule per amine group on the peptide. No unexpected or unusually high safety hazards were encountered.

##### *Synthesis, purification, and characterization of 2AM-L and 2AM-$L_{CC}$.*[1]

*Loading of the resin.* 2-Chlorotrityl chloride resin (300 mg, 1.2 mmol/g) was added to a Bio-Rad Poly-Prep chromatography column (10 mL). The resin was suspended in dry $CH_2Cl_2$ (10 mL) and allowed to swell for 30 min. The solution was drained from the resin and a solution of Fmoc-Gly-OH (0.50 equiv, 53.5 mg, 0.18 mmol) in 6% (v/v) 2,4,6-collidine in dry $CH_2Cl_2$ (8 mL) was added immediately and the suspension was gently agitated for 12 h. The solution was then drained and a mixture of $CH_2Cl_2$/MeOH/*N,N*-diisopropylethylamine (DIPEA) (17:2:1, 10 mL) was added immediately. The mixture was gently agitated for 1 h to cap the unreacted 2-chlorotrityl chloride resin sites. The resin was then washed with dry $CH_2Cl_2$ (2x) and dried by passing nitrogen through the vessel. This procedure typically yields 0.12–0.15 mmol of loaded resin (0.4–0.5 mmol/g loading).

*Peptide coupling.* The Fmoc-Gly-2-chlorotrityl resin generated from the previous step was transferred to a peptide synthesis vessel and submitted to cycles of peptide coupling with Fmoc-protected amino acid building blocks. The linear peptide was synthesized from the *C*-terminus of $G_{25}$ to the *N*-terminus of $S_{26}$ (Scheme 1). Each coupling cycle consisted of i. Fmoc-deprotection with 20% (v/v) piperidine in DMF for 5–10 min, ii. washing with DMF (3x), iii. coupling of the amino acid (0.75 mmol,

5 equiv) in the presence of HCTU (0.675 mmol, 4.5 equiv) and 20% (v/v) *N*-methylmorpholine (NMM) in DMF for 30 min, iv. washing with DMF (3x). The Fmoc-Phe-OH that follows the *N*-methyl phenylalanine was double coupled (0.75 mmol, 5 equiv.) in the presence of HATU and HOAT (0.675 mmol, 4.5 equiv) and allowed to react for 1 h per coupling. After coupling of the last amino acid, the terminal Fmoc group was removed with 20% (v/v) piperidine in DMF. The resin was transferred from the peptide synthesis vessel to a Bio-Rad Poly-Prep chromatography column.

*Cleavage of the peptide from the resin.* The linear peptide was cleaved from the resin by agitating the resin for 1 h with a solution of 1,1,1,3,3,3-hexafluoroisopropanol (HFIP) in $CH_2Cl_2$ (1:4, 7 mL) in the Bio-Rad Poly-Prep column.[2] The suspension was filtered through the frit of the Poly-Prep column and the filtrate was collected in a 250-mL round-bottomed flask. The resin was washed with additional HFIP in $CH_2Cl_2$ (1:4, 7 mL) and then with $CH_2Cl_2$ (2×10 mL). The combined filtrates were concentrated by rotary evaporation to give a white solid. The white solid was further dried by vacuum pump to afford the crude protected linear peptide, which was cyclized without further purification.

*Cyclization of the linear peptide.* The crude protected linear peptide was dissolved in dry DMF (150 mL). HOBt (114 mg, 0.75 mmol, 5 equiv) and HBTU (317 mg, 0.75 mmol, 5 equiv) were added to the solution. DIPEA (0.33 mL, 1.8 mmol, 12 equiv) was added to the solution and the mixture was stirred under nitrogen for 24 h. The mixture was concentrated under reduced pressure to afford the crude protected cyclic peptide.

*Global deprotection of the cyclic peptide.* The protected cyclic peptide was dissolved in TFA/triisopropylsilane (TIPS)/$H_2O$ (18:1:1, 20 mL) in a 250-mL round-bottomed flask equipped with a nitrogen-inlet adaptor. The solution was stirred for 1.5 h. The reaction mixture was then concentrated by rotary evaporation under reduced pressure to afford the crude cyclic peptide as a thin yellow film on the side of the round-bottomed flask. The crude cyclic peptide was immediately subjected to purification by reverse-phase HPLC (RP-HPLC), as described below.

*Reverse-phase HPLC purification.* The crude cyclic peptide was dissolved in $H_2O$ and acetonitrile (7:3, 10 mL), and the solution was filtered through a 0.2 µm syringe filter. The filtrate was injected onto an Agilent Zorbax 300SB-C18 semi-preparative column (21.2 mm x 250 mm, 7 µm particle size) with an Agilent Prep 100Å C18 guard column (21.2 mm x 10 mm) on a Rainin Dynamax HPLC with a flow rate of 20.0 mL/min. The peptides were eluted with a gradient of acetonitrile (20–45% over 90 minutes). Elution was monitored at 214 nm with the accompanying DA Rainin HPLC software. Pure fractions were identified by analytical reverse-phase HPLC using an Agilent 1200 instrument equipped with a Phenomonex Aeris PEPTIDE 2.6u XB-C18 column and then combined and lyophilized to yield pure 2AM-L or 2AM-$L_{CC}$ as fluffy white solids. Syntheses typically yielded 30–40 mg of each peptide as the TFA salt. The lyophilized peptides were analyzed by LC-MS to confirm the purity of the peptide.

***Scheme 1.*** Synthetic scheme for 2AM-L. Synthesis of 2AM-L$_{CC}$ is performed identically, except L$_{17}$ and A$_{21}$ are mutated to cysteine.

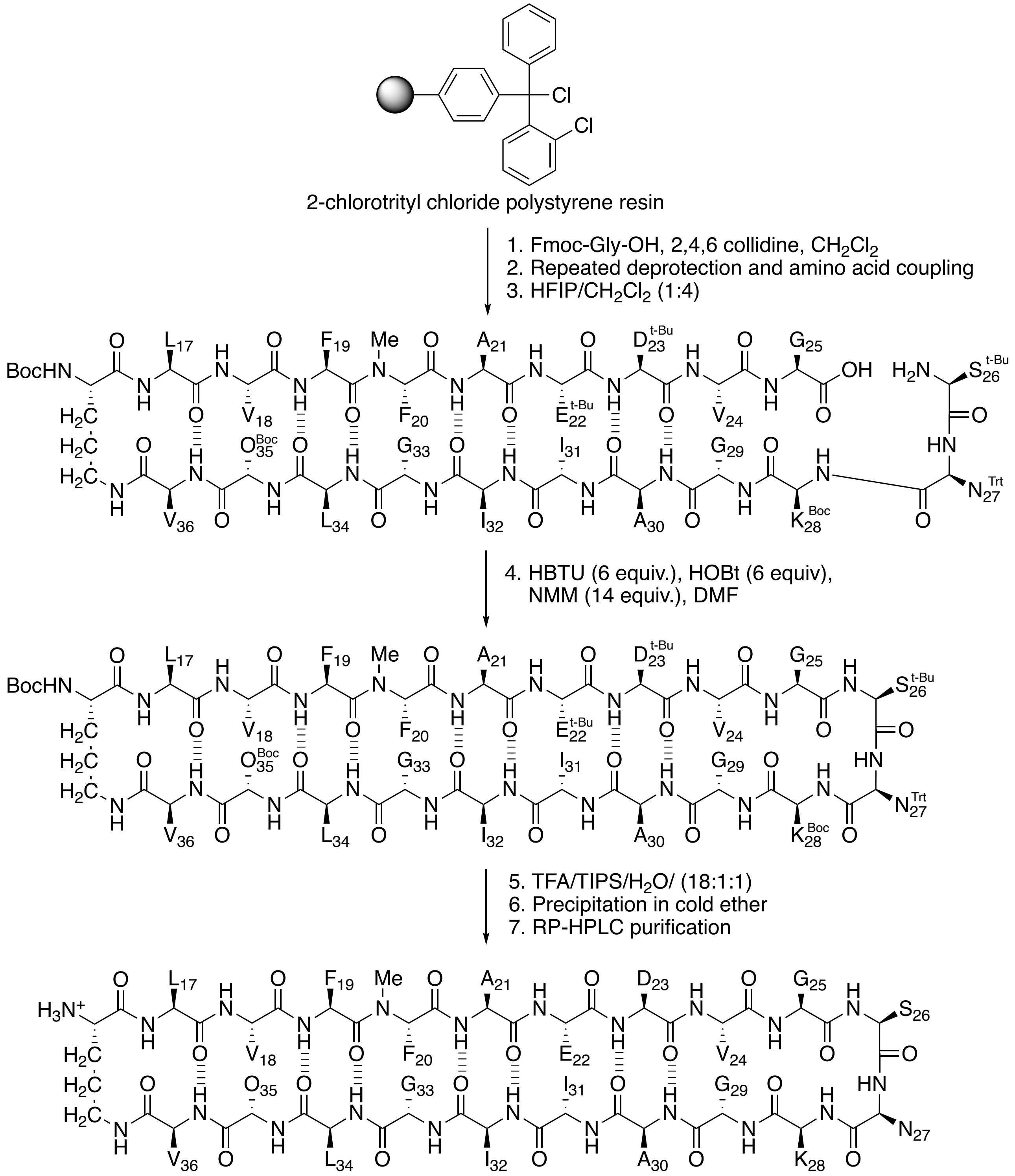


2AM-L (TFA salt)

***Synthesis, purification, and characterization of 2AT-L.***

*Synthesis of 2AT-L.* 2AT-L was synthesized by oxidizing 2AM-$L_{CC}$ in 20% aqueous DMSO with 60 µM triethylamine (TEA).[3,4] A 6 mM solution of lyophilized 2AM-$L_{CC}$ was prepared gravimetrically by dissolving the peptide in an appropriate volume of deionized water. An appropriate volume of DMSO was then added to achieve a final concentration of 20% (v/v) DMSO. An appropriate volume of TEA was then added to achieve a final concentration of 60 µM TEA. In a representative procedure, 30 mg of the 2AM-$L_{CC}$ TFA salt (0.012 mmol) was dissolved in 1.576 mL of deionized water, and then 0.394 mL of DMSO was added, followed by 16.48 µL TEA. The reaction was carried out in a capped 25 mL glass scintillation vial with gentle swirling (80–90 RPM) on a rotating platform at room temperature for 72 h.

*LC-MS analysis of the 2AM-$L_{CC}$ oxidation reaction mixture.* LC-MS analysis of the 2AM-$L_{CC}$ oxidation reaction mixture was performed after 72 h on an ACQUITY UPLC H-class system, Xevo G2-XS QTof (Waters Corp.) equipped with a Protein BEH C4 column (300 Å, 1.7 µm, 2.1 mm X 50 mm, Waters Corp). For LC-MS, the oxidation reaction mixture was diluted 100x by combining 1 µL of the oxidation reaction mixture with 199 µL of water. A 5 µL aliquot of the diluted oxidation reaction mixture was injected onto the column and eluted with gradient of Buffer A consisting of 0.1% formic acid in water (Water LC-MS #9831-02, J.T. Baker; Formic Acid LC-MS #85178, Thermo Scientific) and Buffer B, acetonitrile (Acetonitrile UHPLC-MS #A956, Thermo Scientific). Gradient table listed below:

LC-MS elution gradient table (30-minute method)

| time (min) | flow rate (mL/min) | %A | %B |
|---|---|---|---|
| initial | 0.3 | 97 | 3 |
| 1.0 | 0.3 | 97 | 3 |
| 25.0 | 0.3 | 50 | 50 |
| 27.0 | 0.3 | 10 | 90 |
| 27.5 | 0.3 | 10 | 90 |
| 29.0 | 0.3 | 97 | 3 |
| 30.0 | 0.05 | 97 | 3 |

*Purification of 2AT-L.* The 2AM-$L_{CC}$ oxidation reaction mixture was subjected to an initial reverse-phase LC purification by directly injecting the reaction mixture onto a Biotage® Isolera One flash chromatography instrument equipped with a Biotage® Isolera Sfär Bio C18 D - Duo 300 Å 20 µm 25 g column. The reaction mixture was injected at 20% aqueous $CH_3CN$ and eluted with a gradient of 20–50% $CH_3CN$. Fractions containing 2AT-L were identified by LC-MS as described above and then frozen and lyophilized to afford a white powder containing a mixture of products composed of predominantly 2AT-L. The product mixture was then subjected to reverse-phase HPLC purification by dissolving the white powder in 1–3 mL of 20% aqueous $CH_3CN$ and then injecting the solution onto an Agilent Zorbax 300SB-C18 semi-preparative column (21.2 mm x 250 mm, 7 µm particle size) with an Agilent Prep 100 Å C18 guard column (21.2 mm x 10 mm) on a Rainin Dynamax HPLC with a flow rate of 20.0 mL/min and eluted with a gradient of 20–45% $CH_3CN$ over 90 min. In this HPLC purification, the column was submerged in a 60 ºC water bath heated by a thermal immersion circulator. Fractions containing pure 2AT-L were identified by LC-MS using the 30-minute method described above and then combined, frozen, and lyophilized to afford 2AT-L as a white powder. The powder was analyzed by LC-MS in a similar fashion to that described above to confirm the purity of 2AT-L.

In some instances, a significant number of fractions containing 2AT-L from the HPLC purification were contaminated with intramolecular disulfide monomer. In these instances, the intramolecular disulfide monomer (~2.2 kDa) was removed from 2AT-L (~6.6 kDa) by spin filtration as follows: a solution of the product mixture was prepared in 20 mL of 20% aqueous $CH_3CN$ and then centrifuged through a 3 kDa molecular weight cutoff protein concentrator with a PES filter (Pierce™ catalog # 88526) at 3800 x g until the volume in the upper chamber of the spin filter reached 0.5–1 mL. Multiple rounds of centrifugation were performed (typically 3–6 rounds), replenishing the 20 mL of 20% aqueous $CH_3CN$ at the beginning of each round. After the third round, the solution in the upper chamber was analyzed by LC-MS as described above to determine if all intramolecular disulfide monomer had been removed. If intramolecular disulfide monomer remained, additional rounds of centrifugation were performed. Once all the intramolecular disulfide monomer was removed, the solution in the upper chamber of the spin filter was transferred to a 15 mL conical tube. The reservoir of the upper chamber of the spin filter was rinsed with deionized water and the water rinse was transferred to the 15 mL conical tube. The solution was then frozen in dry ice and lyophilized to yield a fluffy white powder.

Typical syntheses yielded ~5 mg of >95% pure 2AT-L as the TFA salt from a 0.1 mmol scale synthesis of 2AM-$L_{CC}$.

*LC-MS characterization of 2AT-L.* The lyophilized 2AT-L was analyzed by LC-MS to confirm the purity of the trimer. LC-MS analysis was performed on an ACQUITY UPLC H-class system, Xevo G2-XS QTof (Waters Corp.) equipped with a Protein BEH C4 column (300 Å, 1.7 µm, 2.1 mm X 50 mm, Waters Corp). For LC-MS, a 10 mg/mL solution of 2AT-L was prepared gravimetrically by dissolving 1.0 mg of the peptide in 100 µL of deionized water. A 0.05 mg/mL solution of 2AT-L was then created by combining 1 µL of the 10 mg/mL solution with 199 µL of deionized water. The 0.05 mg/mL solution of 2AT-L was then further diluted with deionized water to create a 0.005 mg/mL solution. A 5 µL portion of the 0.005 mg/mL 2AT-L solution was injected onto the column and eluted with gradient of Buffer A consisting of 0.1% formic acid in water (Water LC-MS #9831-02, J.T. Baker; Formic Acid LC-MS #85178, Thermo Scientific) and Buffer B, acetonitrile (Acetonitrile UHPLC-MS #A956, Thermo Scientific). Gradient table listed below:

LC-MS elution gradient table (30-minute method)

| time (min) | flow rate (mL/min) | %A | %B |
|---|---|---|---|
| initial | 0.3 | 97 | 3 |
| 1.0 | 0.3 | 97 | 3 |
| 25.0 | 0.3 | 50 | 50 |
| 27.0 | 0.3 | 10 | 90 |
| 27.5 | 0.3 | 10 | 90 |
| 29.0 | 0.3 | 97 | 3 |
| 30.0 | 0.05 | 97 | 3 |

***X-ray crystallography of 2AT-L.***[1]

*Crystallization procedure for 2AT-L.* Initial crystallization conditions were determined using the hanging-drop vapor-diffusion method. Crystallization conditions were screened for 2AT-L using three crystallization kits in a 96-well plate format (Hampton Index, PEG/Ion, and Crystal Screen). Three 150 nL hanging drops that differed in the ratio of peptide to well solution were made per condition in each 96-well plate for a total of 864 experiments. Hanging drops were made by combining an appropriate volume of 2AT-L (10 mg/mL in 18 MΩ water) with an appropriate volume of well solution to create three 150 nL hanging drops with 1:1, 1:2, and 2:1 2AT-L:well solution. The hanging drops were made using a TTP LabTech Mosquito nanodisperse instrument. Crystals of 2AT-L suitable for X-ray diffraction grew in a solution of 0.1 M Tris at pH 8.3, 0.2 M $MgCl_2$, 2.8 M 1,6-hexanediol.

Crystallization conditions for 2AT-L were optimized using a 4x6 matrix Hampton VDX 24-well plate. The pH of Tris was varied in each row in increments of 0.25 pH units (8.05, 8.30, 8.55, and 8.80) and the 1,6-hexanediol concentration in each column in increments of 0.2 M (2.2 M, 2.4 M, 2.6 M, 2.8 M, 3.0 M, 3.2 M). For the first well in the 4x6 matrix we combined 100 μL of 1 M Tris buffer at pH 8.05, 440 μL of 5 M 1,6-hexanediol solution, 100 μL of 1 M $MgCl_2$, and 360 μL of 18 MΩ water. The other wells were prepared in analogous fashion, by combining 100 μL of Tris buffer of varying pH, 1,6-hexanediol in varying amounts, $MgCl_2$, and 18 MΩ water for a total volume of 1 mL in each well.

Three hanging-drops were prepared per borosilicate glass slide by combining a 10 mg/mL solution of 2AT-L (1 μL) and the well solution (1 μL) in a ratio of 1:1, 2:1, and 1:2. Slides were inverted and pressed firmly against the silicone grease surrounding each well. Crystals of 2AT-L grew in ~96 h. Crystals were harvested with a nylon loop attached to a copper or steel pin and flash frozen in liquid nitrogen prior to data collection.

*X-ray crystallographic data collection, data processing, and structure determination.* Diffraction data for 2AT-L were collected on the Advanced Light Source Synchrotron at the Berkeley Center for Structural Biology on beamline 5.0.1. at 0.97 Å wavelength with 0.5° oscillation. Diffraction data were collected using b4. Diffraction data were scaled and merged using XDS and pointless and aimless. The resolution was limited to the $CC_{1/2}$ resolution of 1.8 Å. Coordinates for the anomalous signal from *para*-iodo-phenylalanine were determined by HySS in the Phenix software suite 1.10.1. Electron density maps were generated using anomalous coordinates determined by HySS as initial positions in Autosol. Molecular manipulations of the model were performed with Coot. Coordinates were refined in the $C222_1$ space group with phenix.refine. The optimal number and composition of the TLS groups were chosen automatically by phenix.refine.[5,6]

***SDS-PAGE and silver staining.***[1]

Running buffers for Tricine SDS-PAGE were prepared according to recipes detailed in Schägger, H. *Nat. Protoc.* 2006, *1*, 16–22.[7] 2AT-L and 2AM-L were run on a 16.5% Mini-PROTEAN® Tris-Tricine Gel (Bio-Rad 4563063) with the Spectra™ Multicolor Low Range Protein Ladder (ThermoFisher Scientific, catalog # 26628).

*Sample preparation and gel running.* Lyophilized 2AT-L and 2AM-L were dissolved in deionized water to a concentration of 10 mg/mL. Aliquots of the 10 mg/mL solutions were then used to create 1.0 mg/mL solutions in 1X SDS-PAGE loading buffer (112.5 mM Tris buffer (pH 8.0) with 2% (w/v) SDS and 6% (v/v) glycerol). The 1.0 mg/mL solutions were then serially diluted in 1X SDS-PAGE loading buffer to create 0.5, 0.25, and 0.125 mg/mL solutions. A 5.0-μL aliquot of each dilution was run on the gel. Once the gels were loaded, the gel-running apparatus was moved to a 4 °C cold room and allowed to equilibrate to 4 °C for 30 min. After equilibration, the gels were run at a constant 30 volts until adequate band separation was observed for the protein ladders.

*Silver staining.* Staining with silver nitrate was used to visualize 2AT-L and 2AM-L in the gels. Reagents for silver staining were prepared according to procedures detailed in Simpson, R. J. *CSH Protoc.* 2007.[8] [The sodium thiosulfate solution, silver nitrate solution, and developing solution were prepared fresh each time silver staining was performed. Furthermore, the purity of the sodium carbonate in the developing solution was >99.5%].

For silver staining, the gel was removed from the cast and submerged and rocked in fixing solution (50% (v/v) methanol and 5% (v/v) acetic acid in deionized water) for 20 min. The fixing solution was then discarded and the gel was rocked in 50% (v/v) aqueous methanol for 10 min. The 50% methanol was then discarded and the gel was rocked in deionized water for 10 min. The water was then discarded and the gel was soaked in 0.02% (w/v) sodium thiosulfate in deionized water for 1 min. The sodium thiosulfate was then discarded and the gel was rinsed with deionized water for 1 min (2X). After the last rinse, the gel was submerged in chilled 0.1% (w/v) silver nitrate in deionized water and rocked at 4 °C for 20 min. The silver nitrate solution was then discarded and the gel was rinsed with deionized water for 1 min (2X). To develop the gel, the gel was incubated in developing solution (2% (w/v) sodium carbonate, 0.04% (w/v) formaldehyde until the desired intensity of staining was reached (~1–3 min). When the desired intensity of staining was reached, the development was stopped by discarding the developing solution and submerging the gel in 5% aqueous acetic acid. The silver-stained gels were immediately visualized with a ChemiDoc™ MP Imaging System using the "Optimal Auto-exposure" exposure under the "Silver Stain Gel" application of the Image Lab Touch Software.

***Size exclusion chromatography (SEC).***[1]

2AT-L and 2AM-L were studied by size exclusion chromatography (SEC) in TBS (50 mM Tris buffer (pH 7.5) and 150 mM NaCl) as follows: Each compound was dissolved in deionized water to a concentration of 10 mg/mL. The 10-mg/mL solutions were then diluted to 1 mg/mL by adding 80 μL of the 10-mg/mL solutions to 720 μL of TBS. The 1-mg/mL solutions were centrifuged at 13,500 RPM for 30 seconds to remove precipitate, and the supernatant was and then loaded onto a GE Superdex 75 10/300 GL column at 0.5 mL/min over 1 min. After loading, the samples were eluted with TBS at 1 mL/min. Chromatograms were recorded at 214 nm and normalized to the highest absorbance value. The standards

Dextran blue (2000 kDa), cytochrome C (12.4 kDa), aprotinin (6.5 kDa), and vitamin $B_{12}$ (1.3 kDa) were run in the same fashion.

***Dynamic light scattering (DLS).***[1]

Dynamic light scattering was measured using a Malvern Zetasizer ZS Nano DLS at ambient temperature (ca. 20ºC). Solutions of 2AT-L and 2AM-L (25 μM) were prepared in 10 mM potassium phosphate buffer at pH 7.4, centrifuged at 17,000 x g for 2 minutes, and then the supernatants were transferred to 1 cm disposable plastic cuvettes. Data were collected in 10 seconds time intervals and averaged over 3 measurements. The scattering was measured with a 173º backscattering angle.

***Circular dichroism (CD) spectroscopy.***[1]

A 50-μM solution of either 2AT-L or 2AM-L was prepared by diluting an aliquot from a 10 mg/mL stock solution of each compound prepared in deionized water with 10 mM potassium phosphate buffer at pH 7.4. Each solution was transferred to a 1 mm quartz cuvette for data acquisition. CD spectra were acquired on a Jasco J-810 circular dichroism spectropolarimeter at room temperature. Data were collected using 0.2 nm intervals from 260 nm to 190 nm and averaged over five accumulations with smoothing.

***Cell-based toxicity assays of 2AT and KLT in SH-SY5Y cells.***[1]

The cellular toxicity of 2AT-L and 2AM-L was assessed by measuring LDH release, ATP levels, and caspase-3/7 activation in SH-SY5Y cells exposed to a twofold dilution series (50–1.6 μM, 0 μM) of 2AT-L or 2AM-L for 72 h. LDH release, ATP levels, and caspase-3/7 activation were measured using the following assays:

- LDH release: CyQUANT™ LDH Cytotoxicity Assay (ThermoFisher Scientific; cat# C20301)
- ATP levels: CellTiter-Glo® 2.0 Cell Viability Assay (Promega Corporation; cat. # G9242).

- Caspase-3/7 activation: Apo-ONE® Homogeneous Caspase-3/7 Assay (Promega Corporation; cat. # G7790).

Serial dilutions of 2AT-L and 2AM-L were first prepared in a replica 96-well plate and then transferred to the plate containing cells. Three technical replicates were performed for each experiment.

*Preparation of SH-SY5Y cells for the toxicity assays.* SH-SY5Y cells were plated in the inner 60 wells (rows B–G, columns 2–10) of cell culture-treated, black-walled, half-area, flat-bottom, clear-bottom 96-well plates (Corning™ cat. # 3882) at 30,000 cells/well. DMEM:F12 media (100 µL) was added to the outer wells (rows A and H and columns 1 and 12), to create an evaporative barrier and ensure the greatest reproducibility of data generated from the inner wells. The cells were plated in 50 µL of a 1:1 mixture of DMEM:F12 media supplemented with 10% fetal bovine serum, 100 U/mL penicillin, and 100 µg/mL streptomycin and incubated at 37 °C in a 5% $CO_2$ atmosphere for 24 hours to allow the cells to adhere to the bottoms of the wells.

*Preparation of 2AT-L and 2AM-L for toxicity assays.* 10 mg/mL stock solutions of the 2AT-L and 2AM-L TFA salts were prepared gravimetrically by dissolving 1.0 mg of each compound in 100 µL of deionized water that had been passed through a 0.2 µm filter. The 10 mg/mL solution of the 2AT-L TFA salt is equivalent to 1.25 mM 2AT-L. The 10 mg/mL solution of the 2AM-L TFA salt is equivalent to 3.75 mM 2AM-L. The 10 mg/mL stock solutions were prepared in a 1.7 mL microcentrifuge tube and stored at -20 ºC when not in use.

After the cells had adhered to the bottoms of the wells, the 10 mg/mL stock solutions of each compound were used to create a replica 96-well plate (Corning™ cat. # 353075) containing a twofold dilution series (50–1.6 µM, 0 µM) of 2AT-L and 2AM-L as follows: (1) Add 75 µL of serum-free, Phenol Red-free DMEM:F12 media to the wells in columns 3–11 of rows B–G of the 96-well plate. (2) In the wells of column 2 of rows B–D, prepare 150 µL of a 50 µM solution of 2AT-L by adding 6.0 µL of the 1.25 mM stock solution of 2AT-L to 144.0 µL of serum-free, Phenol Red-free DMEM:F12 media; in the wells of column 2 of rows E–G prepare 150 µL of a 50 µM solution of 2AM-L by adding 2 µL of the 3.75

mM stock solution of 2AM-L to 148 µL of serum-free, Phenol Red-free DMEM:F12 media. (3) using a multi-channel pipette, perform a twofold serial dilution of the trimers by transferring 75 µL of the 50 µM solutions from column 2 to column 3 and mixing by pipetting up and down 8–10 times, and then transferring 75 µL from column 3 to column 4 and mixing, and so on, ending the serial dilution on column 7, with column 8 not receiving any trimer and thus constituting the 0-µM vehicle control.

*Treatment of the SH-SY5Y cells with 2AT-L and 2AM-L.* A multi-channel pipette was used to remove the media from the wells of the 96-well plate containing cells. A multi-channel pipette was then used to immediately add 50 µL from each well of the 96-well replica plate containing 2AT-L and 2AM-L dilution series to each respective well of the 96-well plate containing cells. The plate was incubated at 37 °C in a 5% $CO_2$ atmosphere for 72 hours and then the assays were performed according to manufacturer's instructions.

*CyQUANT™ LDH Cytotoxicity Assay.* After 72 hours, LDH release was measured using a CyQUANT™ LDH Cytotoxicity Assay according to the manufacturer's instructions, except the volumes of assay reagent added to the wells were halved, to accommodate the half-area wells. A 50 µL aliquot of the supernatant media from each well was transferred to a new 96-well plate and 50 µL of LDH substrate solution, prepared according to manufacturer's protocol, was added to each well. The treated plates were stored in the dark for 30 min. The absorbance of each well was measured at 490 nm on a ThermoFisher Scientific Varioskan Lux plate reader. The absorbance measurements for the three replicates of each treatment group were averaged and the standard deviations were calculated using GraphPad Prism. The data were then plotted using GraphPad Prism.

*CellTiter-Glo® 2.0 Cell Viability Assay.* After 72 hours, ATP levels were measured using a CellTiter-Glo® 2.0 Cell Viability Assay according to the manufacturer's instructions, except the volumes of assay reagent added to the wells were halved, to accommodate the half-area wells. The 96-well plate was removed from the incubator and allowed to come to room temperature for 30 minutes. Once the plate reached room temperature, a 4 mL aliquot of CellTiter-Glo® 2.0 reagent was thawed and then added to a reagent reservoir (Thermo Scientific cat. # 8093-11). A multi-channel pipette was then used to transfer 50 μL of the CellTiter-Glo® 2.0 reagent to each well containing cells on the 96-well plate. The plate was then shaken on a rotating shaker for 2 minutes at 100 RPM. The luminescence from each well was then measured on a Promega GloMax® Discover Microplate Reader. The luminescence readings for the three replicates of each treatment group were averaged and the standard deviations were calculated using GraphPad Prism. The data were then plotted using GraphPad Prism.

*Apo-ONE® Homogeneous Caspase-3/7 Assay.* After 72 hours, caspase-3/7 activation was measured using an Apo-ONE® Homogeneous Caspase-3/7 Assay according to the manufacturer's instructions, except the volumes of assay reagent added to the wells were halved, to accommodate the half-area wells. The Apo-ONE® Homogeneous Caspase-3/7 Assay reagents were prepared according to the instructions. The 96-well plate was removed from the incubator and the compound-containing media was removed and replaced with 25 μL of fresh serum-free DMEM/F12 media. A 25-μL aliquot of assay reagent was then added to each well. The plate was sealed with a clear adhesive plate sealer and fluorescence was monitored over 18 h while shaking at 250 rpm using a ThermoFisher Scientific Varioskan Lux plate reader (excitation at 499 nm, emission at 521 nm). The end-point fluorescence readings for the three replicates of each treatment group were averaged and the standard deviations were calculated using GraphPad Prism. The data were then plotted using GraphPad Prism.

***Generation of pAb$_{2AT\text{-}L}$.***

*Rabbit immunization.* pAb$_{2AT\text{-}L}$ was generated by Pacific Immunology (Ramona, CA, www.pacificimmunology.com) using standard custom antibody production procedures. Briefly, 2AT-L was conjugated to the carrier protein keyhole limpet hemocyanin (KLH) using standard EDC conjugation chemistry. The EDC was obtained from Thermo Scientific (catalog # 22980) and the conjugation was performed according to manufacturer's instructions. Two New Zealand white rabbits were then immunized with the 2AT-L/KLH conjugate in complete Freunds adjuvant. The rabbits were boosted after ~21 days with the 2AT-L/KLH conjugate in incomplete Freunds adjuvant, and then boosted again ~21 days later. Production bleeds were performed 7 and 21 days after the second boost to yield ~25 mL of antisera from each rabbit. The rabbits were then boosted on a monthly schedule, with production bleeds occurring 7 and 21 days after each boost.

*Affinity purification of pAb$_{2AT\text{-}L}$.* pAb$_{2AT\text{-}L}$ was purified from the antisera using affinity chromatography. To create the affinity chromatography column, 10 mg of the 2AT-L TFA salt was dissolved in 2 mL DMSO. The 2AT-L/DMSO solution was then added to 10 mL PBS (pH 7.4) to create a 1 mg/mL 2AT-L solution in PBS with 20% DMSO. The 1 mg/mL 2AT-L solution was then added to 750 mg dry NHS-activated agarose resin (Thermo Scientific, catalog # 26197) in a 15 mL polypropylene conical tube. The 2AT-L/agarose suspension was then continuously inverted using a tube rotator for two hours at room temperature. After the two-hour incubation, the agarose was transferred to a Bio-Rad Poly-Prep chromatography column (10 mL) and washed 3x with PBS. The agarose was then incubated in 5 mL 1 M Tris buffer (pH 8.5) for 1 hour at room temperature on a tube rotator to cap unreacted NHS-ester groups, and then washed 3x with PBS.

For the affinity purification of pAb$_{2AT\text{-}L}$ from the antisera, the 2AT-L-agarose was transferred to a 50 mL CrystalCruz® glass chromatography column. The antisera from a single production bleed from both rabbits were combined and then filtered through 0.45 μm syringe filters. The filtered antisera were then added to the column containing the 2AT-L-agarose and rocked on a rocker for ~3 hours at room temperature or overnight at 4 ºC. Next, the antisera were drained from the column into a 50 mL polypropylene conical tube, and the 2AT-L-agarose was washed with ice-cold PBS. Washing was performed with 50 mL portions of ice-cold PBS until the absorbance at 214 nm of the wash buffer eluent measured less than 0.05 on a NanoDrop One/One$^c$ Microvolume UV-Vis Spectrophotometer.

pAb$_{2AT\text{-}L}$ was then eluted from the 2AT-L-agarose using ice-cold 0.2 M glycine buffer (pH 1.85). The pAb$_{2AT\text{-}L}$ elution was performed by adding the glycine buffer in 1 mL portions and then collecting the eluent into 1.7 mL microcentrifuge tubes containing 0.5 mL 1 M Tris buffer (pH 8.5). Typically, 10–15 1-mL portions of the glycine buffer were sufficient to elute all of $pAb_{2AT-L}$ from the 2AT-L-agarose. The eluents were then combined and transferred to a 30 kDa molecular weight cutoff filter (30K MWCOF) (Thermo Scientific catalog # 88531) and buffer exchange with PBS was performed. The 30K MWCOF was centrifuged in a swinging bucket centrifuge at 3800 x g until the volume fell below 1 mL, after which the buffer was replenished with ice-cold PBS up to the 20 mL marker on the 30K MWCOF. This process was repeated at least six times. For the final centrifugation step, the $pAb_{2AT-L}$ volume was allowed to reach ~0.5 mL in the 30K MWCOF and was then transferred to a 1.7 mL microcentrifuge tube. The $pAb_{2AT-L}$ concentration was determined using a BCA assay and then adjusted to 1 mg/mL with PBS. Typical affinity purifications yielded 1–3 mg of $pAb_{2AT-L}$ from 50 mL of antisera. The affinity purified $pAb_{2AT-L}$ was then portioned into 50 µL aliquots, which were stored at -80 ºC and thawed as needed. Once thawed, the aliquot was stored at 4 ºC.

***Indirect ELISA of $pAb_{2AT-L}$ against 2AT-L, 2AM-L, and BSA.***

Indirect ELISA was used to determine the selectivity of $pAb_{2AT-L}$ for 2AT-L and 2AM-L, as well as the negative control protein bovine serum albumin. Three technical replicates were performed for each experiment. Each aspiration and washing step in the ELISA procedure was performed using a Fisherbrand™ accuWash™ Microplate Washer (catalog # 14-377-577).

*Coating the wells of the ELISA plate with 2AT-L, 2AM-L, and BSA.* 10 mg/mL stock solutions of the 2AT-L and 2AM-L TFA salts were prepared gravimetrically by dissolving 1.0 mg of each compound in 100 µL of deionized water that had been passed through a 0.2 µm filter. 1 µg/mL solutions of 2AT-L and 2AM-L were then prepared by adding 1 µL of the 10 mg/mL stocks to 10 mL carbonate buffer (15 mM $Na_2CO_3$, 35 mM $NaHCO_3$, 0.02% (w/v) sodium azide, pH 9.5). The 1 µg/mL solutions of 2AT-L and 2AM-L were then poured into a reagent reservoir and a multichannel pipette was used to transfer 50 µL to the appropriate wells of a Thermo Scientific™ Maxisorp 96-well plate (catalog# 12-565-135). Each well contained a total of 50 ng 2AT-L or 2AM-L. 2AT-L was added to all wells in columns 1–3 and 2AM-L was added to all wells in columns 4–6. For BSA, a 1% (w/v) solution of BSA was prepared by dissolving 100 mg of BSA (Fraction V) (Fisher BioReagents™, catalog # BP1600-100) in 10 mL carbonate buffer. 50 μL of the 1% BSA solution was then added to all wells in columns 7–9. The 96-well plate was then sealed with an adhesive 96-well plate seal (Axygen, catalog# PCR-SP) and incubated overnight (~16 hours) at room temperature on a rotating shaker set to 90 RPM.

*Treating the ELISA plate with $pAb_{2AT\text{-}L}$.* The next day, 20 mL of 1% BSA was prepared by adding 200 mg of BSA (Fraction V) to 20 mL PBS. The solutions of 2AT-L, 2AM-L, and BSA were aspirated from the wells of the 96-well plate and then washed 1x with PBST (10 mM $Na_2HPO_4$, 1.8 mM $KH_2PO_4$, 137 mM NaCl, 2.7 mM KCl, 0.5% Tween-20). Next, a multichannel pipette was used to transfer 75 μL of 1% BSA to each well, the plate was sealed, and incubated for at least 1 hour at room temperature on a rotating shaker set to 90 RPM to block uncoated sites in the wells. During the last 15 minutes of the blocking step, a 1 μg/mL $pAb_{2AT\text{-}L}$ solution was prepared by combining 1 μL of the 1 mg/mL affinity purified $pAb_{2AT\text{-}L}$ with 999 μL of 1% BSA. When the blocking step was complete, the wells were aspirated and washed 3x with PBST. Using a multi-channel pipette, 50 μL of 1% BSA was then added to the wells in the bottom seven rows of the 96-well plate (rows B–H). 75 μL of the 1 μg/mL $pAb_{2AT\text{-}L}$ solution was then added to the wells in the top row (row A). A multi-channel pipette was then used to create a three-fold dilution series of $pAb_{2AT\text{-}L}$ by transferring 25 μL of the 1 μg/mL $pAb_{2AT\text{-}L}$ solution from row A to row B and then mixing up and down 8 times, and then 25 μL was transferred from row B to row C, and so on, until the last 25 μL had been transferred to row H. The 96-well plated was then sealed with an adhesive plate seal and incubated for 2 hours at room temperature on a rotating shaker set to 90 RPM.

*Treating the ELISA plate with the secondary antibody.* After the 2-hour incubation, the $pAb_{2AT\text{-}L}$ solutions were aspirated and the wells were washed 3x with PBST. A 50 μL portion of AffiPure Goat Anti-Rabbit IgG (H+L) conjugated to horse radish peroxidase (GαR-HRP; Jackson ImmunoResearch, catalog #

111-035-144) diluted 1:10,000 in 1% BSA was then added to each well. The 96-well plated was then sealed with an adhesive plate seal and incubated for 1 hour at room temperature on a rotating shaker set to 90 RPM.

*Developing the ELISA plate.* After the 1-hour incubation, the GαR-HRP solution was aspirated and the wells were washed 3x with PBST. A 50 μL portion of 3,3',5,5'-tetramethybenzidine (TMB) (Millipore Sigma, catalog # ES001-500ML) was then added to each well and allowed to react until the blue color reached a sufficient hue. A 50 μL portion of 1 M aqueous HCl was then added to each well to quench the reaction and the absorbance was measured at 450 nm using a MultiSkan GO plate reader. The absorbance readings for the three replicates were averaged and the standard deviations were calculated using GraphPad Prism. The data were then plotted and fit using GraphPad Prism to estimate the $EC_{50}$ values for 2AT-L and 2AM-L (Figure S2).

***Immunostaining and fluorescence microscopy of human and 5xFAD mouse brain slices.***

*Preparing the human brain tissues for immunostaining.* The LOAD, DSAD, and CAA brain tissues used in this project were provided by the University of California Alzheimer's Disease Research Center (UCI-ADRC) and the Institute for Memory Impairments and Neurological Disorders. The human brain tissues were received as blocks of tissue fixed in 4% paraformaldehyde and stored in PBS with 0.02% sodium azide ($NaN_3$). Upon receipt, the tissue blocks were cryopreserved by soaking the tissue in 15% (w/v) sucrose in PBS with 0.02% $NaN_3$ in a 50 mL conical tube at 4 ºC until the tissue block sunk to the bottom of the tube, and then in 30% (w/v) sucrose in PBS containing 0.02% $NaN_3$ in a similar fashion until the tissue block sunk to the bottom of the tube. The tissue blocks were stored at 4 ºC in the 30% sucrose solution until sectioned. For sectioning, the tissue blocks were mounted and frozen in dry ice on the stage of an Epredia™ HM 430 Sliding Microtome (Fisher, catalog # 22-050-855), and then cut into slices with a thickness of 40 μm. The slices were stored at 4 ºC in a 6-well or 12-well dish in PBS with 0.02% $NaN_3$ until use.

*Preparing the 5xFAD mouse brain tissues for immunostaining.* 5xFAD mice were bred and genotyped by the Chao Family Comprehensive Cancer Center Transgenic Mouse Facility Shared Resource.

The mice were euthanized by intraperitoneal administration of a lethal dose of EUTHASOL® Euthanasia Solution (pentobarbital sodium and phenytoin sodium). The euthanized mice were transcardiac perfused with ice-cold PBS and their brains were then removed. Using a spatula, the excised brains were cut along the deep longitudinal fissure to separate the two hemispheres. One hemisphere was placed in a 1.7 mL microcentrifuge tube, flash frozen in liquid nitrogen, and then stored at -80 ºC for subsequent biochemical studies. The other hemisphere was drop-fixed in 20 mL of 4% paraformaldehyde in a 20 mL glass scintillation vial and fixed for 96 hours at room temperature for the subsequent immunostaining studies. After fixation, the brain hemisphere was washed 5x with PBS containing 0.02% $NaN_3$ and then stored in the PBS/azide solution at 4 ºC until use. To prepare the fixed brain hemisphere for sectioning, the tissue was cryopreserved by soaking the tissue in 15% (w/v) sucrose in PBS with 0.02% $NaN_3$ in the 20 mL glass scintillation vial at 4 ºC until the tissue sunk to the bottom of the tube, and then in 30% (w/v) sucrose in PBS with 0.02% $NaN_3$ in a similar fashion until the tissue sunk to the bottom of the tube. The brain hemispheres were stored at 4 ºC in the 30% sucrose solution until sectioned. For sectioning, the brain hemispheres were mounted and frozen in dry ice on the stage of an Epredia™ HM 430 Sliding Microtome (Fisher catalog # 22-050-855), and then cut into coronal slices with a thickness of 40 µm. The slices were stored at 4 ºC in a 12-well or 24-well dish in PBS with 0.02% $NaN_3$ until use.

*Staining and imaging the human and 5xFAD mouse brain tissue slices.* The human and mouse brain tissue slices were immunostained with $pAb_{2AT-L}$ in a nearly identical manner. All washing, soaking, and incubation steps were performed at room temperature on a rotating platform set to 70 RPM. In some instances, the tissue slices were also stained with the anti-Aβ antibody 6E10, or the amyloid-staining dyes AmyTracker480 (AT480) and AmyTracker680 (AT680).

antibody and dye information for immunostaining

| antibody | host animal | source | working concentration |
|---|---|---|---|
| $pAb_{2AT-L}$ | rabbit | custom made | 3 μg/mL |
| 6E10 | mouse | Biolegend (catalog # 803015) | 3 μg/mL |
| Goat anti-Rabbit IgG Alexa Fluor™ Plus 488 | goat | Invitrogen (catalog # A32731) | 2 μg/mL |
| Goat anti-Mouse IgG Alexa Fluor™ Plus 647 | goat | Invitrogen (catalog # A32728) | 2 μg/mL |

| dye | source | dilution |
|---|---|---|
| AmyTracker480 (AT480) | Ebba Biotech (catalog # A680) | 1:1000 |
| AmyTracker680 (AT680) | Ebba Biotech (catalog # A480) | 1:1000 |
| DAPI | Thermo Scientific™ (catalog # 62248) | 1:1000 |

To begin the immunostaining procedure, the tissue slices were transferred from the storage plates to the wells of clean 12- or 24-well plates containing TBS (50 mM Tris buffer pH 7.4, 150 mM NaCl). The tissue slices were then washed 3x with TBS for 5 minutes. Antigen retrieval was then performed by soaking the tissues in 88% aqueous formic acid for 5 minutes, followed by washing the tissue 3x with TBS for 5 minutes. Permeabilization was then performed by incubating the tissues in TBSA (TBS with 0.1% Triton-X) for 30 minutes. The tissue slices were then blocked by incubating the tissues in TBSB (TBSA with 2% bovine serum albumin Fraction V) for 1 hour. In instances where mouse tissue slices were stained with 6E10, which has a mouse IgG1 isotype, M.O.M. ® (Mouse on Mouse) Blocking Reagent (vector laboratories, catalog # MKB-2213-1) was added to the TBSB used in the blocking step. The blocking solution was then removed and replaced with a primary antibody solution prepared in TBSB. The primary antibodies were used at the concentrations listed in the table above. The tissue slices were incubated in the primary antibody solution overnight (~16 hours). The next day, the primary antibody solutions were removed and the tissue slices were washed 3x with TBSA for 5 minutes. The tissue slices were then re-blocked by incubating the tissues in TBSB containing 5% normal goat serum (Gibco™, catalog # PCN5000) for 30 minutes. The blocking solution was then removed and replaced with an appropriate secondary antibody solution prepared in TBSB with 5% normal goat serum. The secondary antibodies were used at the dilutions listed in the table above.

The tissue slices were incubated in the secondary antibody solution for 2–3 hours. The secondary antibody solutions were then removed and the tissue slices were washed 3x with TBS for 5 minutes. In instances where the AmyTracker dyes were used, the tissue slices were then incubated in a solution of AmyTracker dye prepared in TBS at a 1:1000 dilution for 30 minutes. The dye solution was then removed, and the tissue slices were washed 2x in TBS for 2 minutes. In instances where DAPI was used, the tissue slices were then incubated in a solution of DAPI prepared in TBS at a 1:1000 dilution for 30 minutes. The tissue slices were then washed 3x with TBS for 5 minutes and then mounted on glass slides and cover-slipped. The tissue slices were either imaged immediately or were placed in a slide storage box and stored at 4 ºC until imaged. The stained brain slices were imaged using a Zeiss LSM 900 with Airyscan 2 confocal microscope, or a Keyence BZ-X800 fluorescent microscope. The images in Figures 3, 4, and 5 were acquired on the Zeiss LSM 900 confocal microscope using a 10x, 20x, or 63x objective; the images in Figures S3–S9 were acquired on the BZ-X800 fluorescent microscope using a 4x or 10x objective.

***Preparation of 5xFAD brain protein extracts.***

The protocol for isolating $pAb_{2AT-L}$-reactive species was adapted from a protocol first described by Ashe and co-workers. Frozen brain hemispheres from 8-month-old 5xFAD and wild type mice, isolated and stored as described above, were thawed on ice. A 250 µL portion of ice-cold TBSE (50 mM Tris buffer pH 7.4, 150 mM NaCl, and 1 mM EDTA with Halt™ Protease Inhibitor Cocktail (Thermo Scientific, catalog # 78430)) was then added to the thawed brain tissue. The tissue was then passed through a 20-gauge needle attached to a 1 mL syringe 8 times to mechanically homogenize the tissue. The tissue homogenate was then centrifuged at 17,000 x g in a tabletop centrifuge at 4 ºC for 90 minutes. After centrifugation, the TBSE-soluble protein supernatant was collected in a new 1.7 mL microcentrifuge tube and stored at -20 ºC overnight. A 300 µL portion of ice-cold TBSEd (TBSE with 0.5% Triton-X, 3% (w/v) SDS, and 1% (w/v) sodium deoxycholate) was then added to the tissue pellet. The tissue pellet was then passed through a 1 mL pipette tip attached to a pipette until the suspension smoothly moved through the pipette tip (about 10 x). The tissue pellet homogenate was then centrifuged at 17,000 x g in a tabletop centrifuge at 4 ºC for 90 minutes. After centrifugation, the TBSEd-soluble protein supernatant was collected in a new 1.7 mL microcentrifuge tube and stored at -20 ºC overnight.

The next day, the TBSE and TBSEd protein extracts were removed from the -20 ºC freezer and thawed on ice. The extracts were then centrifuged at 17,000 x g in a tabletop centrifuge at 4 ºC for 90 minutes to remove any precipitate. During centrifugation, two 50 μL portions of Protein A/G Magnetic Beads (Pierce, catalog # 88802) were washed 5 x with TBS. The extracts were then added to the washed Protein A/G beads and incubated on a tube rotator at 4 ºC for 1 hour to immunodeplete the extracts. The Protein A/G beads were then removed and the extracts were transferred to new 1.7 mL microcentrifuge tubes. The total protein concentration in each extract was then quantified using a Pierce™ BCA Protein Assay Kit (Thermo Scientific™, catalog # 23225). The extracts were then aliquoted into portions containing 250 μg total protein and stored at -80 ºC until use.

***Dot blot assay of pAb$_{2AT-L}$ against 5xFAD brain protein extracts.***

The 5xFAD and wild type brain protein extracts were removed from the -80 ºC freezer and thawed on ice. The concentration of each extract was adjusted to 1 mg/mL with ice-cold TBSE or TBSEd. A twofold dilution series (1–0.13 mg/mL) for each extract was then created by diluting the 1 mg/mL protein extracts with ice-cold TBSE or TBSEd. A 1 μL portion from each concentration in the dilution series for each extract was then spotted onto a nitrocellulose membrane, and the membrane was allowed to air-dry for 10 minutes. Three identical membranes were created: one to be probed with pAb$_{2AT-L}$, one to be probed with 6E10, and one to be probed with goat anti-rabbit IgG HRP. Non-reactive sites were blocked on each membrane by rocking the membranes in 5% (w/v) non-fat powdered milk in low-Tween TBS (TBS-lT: 50 mM Tris buffer pH 7.4, 150 mM NaCl, 0.01% Tween 20) for 1 h at room temperature. The membranes were then incubated while rocking overnight at 4 °C in pAb$_{2AT-L}$ (3 μg/mL) or 6E10 (3 μg/mL) prepared in 5% milk solution. The third blot was incubated in a 5% milk solution containing no primary antibody and would serve as the goat anti-rabbit "secondary only" negative control.

The next day, the membranes were washed with 3x TBS-lT for 5 minutes. The $pAb_{2AT-L}$ membrane and the "secondary only" membrane were then treated with a solution of goat anti-rabbit IgG HRP antibody (0.8 μg/mL) (Jackson ImmunoResearch catalog# 111-035-003); the 6E10 membrane was treated with a solution of goat anti-mouse IgG HRP antibody (0.8 μg/mL) (Jackson ImmunoResearch catalog# 115-035-062) The membranes were then washed 3x with TBS-lT for 5 min. A 20 mL portion of chemiluminescent HRP substrate (Thermo Scientific SuperSignal™ West Pico PLUS Chemiluminescent Substrate, catalog # 34580) was prepared according to the manufacturer's instructions and then 5 mL of the prepared substrate was added to each membrane. The membranes were incubated in the HRP substrate for ~5 min before imaging. The blot was imaged using a Bio-Rad GelDoc Go Gel Imaging System using the "Auto-expose" option of the Bio-Rad Image Lab Touch Software.

***Immunoprecipitation LC-MS on 5xFAD brain protein extracts.***

Two 250 μg 5xFAD brain protein extract aliquots were removed from the -80 ºC freezer and thawed on ice. The volume of each brain protein extract was then adjusted to 1 mL with ice-cold TBS. The brain protein extracts were then immunodepleted once more by adding 50 μL of washed Protein A/G magnetic beads to each extract and then incubating the samples at 4 ºC on a tube rotator for 1 hour. The Protein A/G magnetic beads were then removed and 5 μg of $pAb_{2AT-L}$ was added to one of the extracts. No antibody was added to the other extract, as it will serve as the (-) $pAb_{2AT-L}$ control. A 50 μL portion of washed Protein A/G magnetic beads was then added to each extract and the samples were incubated over night at 4 ºC on a tube rotator. The next day, the magnetic beads were separated from the extracts using a magnet and washed 5x with ice cold TBS. The bound contents were eluted from the Protein A/G magnetic beads by incubating the beads in 50 μL 88% aqueous formic acid at room temperature for 5 minutes. The formic acid was then evaporated on a SpeedVac centrifuge and the remaining material was redissolved in

20 μL 20% aqueous acetonitrile containing 0.1% formic acid. The redissolved material was then analyzed by LC-MS. LC-MS was performed on an ACQUITY UPLC H-class system, Xevo G2-XS QTof (Waters Corp.) equipped with a Protein BEH C4 column (300 Å, 1.7 μm, 2.1 mm X 50 mm, Waters Corp). For LC-MS, a 5 μL portion of the redissolved material was injected onto the column and eluted with gradient of Buffer A consisting of 0.1% formic acid in water (Water LC-MS #9831-02, J.T. Baker; Formic Acid LC-MS #85178, Thermo Scientific) and Buffer B, acetonitrile (Acetonitrile UHPLC/MS #A956, Thermo Scientific). Gradient table listed below:

LC/MS elution gradient table (5 minute method)

| time (min) | flow rate (mL/min) | %A | %B |
|---|---|---|---|
| initial | 0.3 | 97 | 3 |
| 0.5 | 0.3 | 97 | 3 |
| 2 | 0.3 | 40 | 60 |
| 2.5 | 0.3 | 40 | 60 |
| 3 | 0.3 | 10 | 90 |
| 3.5 | 0.3 | 10 | 90 |
| 4 | 0.3 | 97 | 3 |
| 5 | 0.3 | 97 | 3 |

### Characterization Data

*Characterization of 2AM-L*

LC-MS of 2AM-L

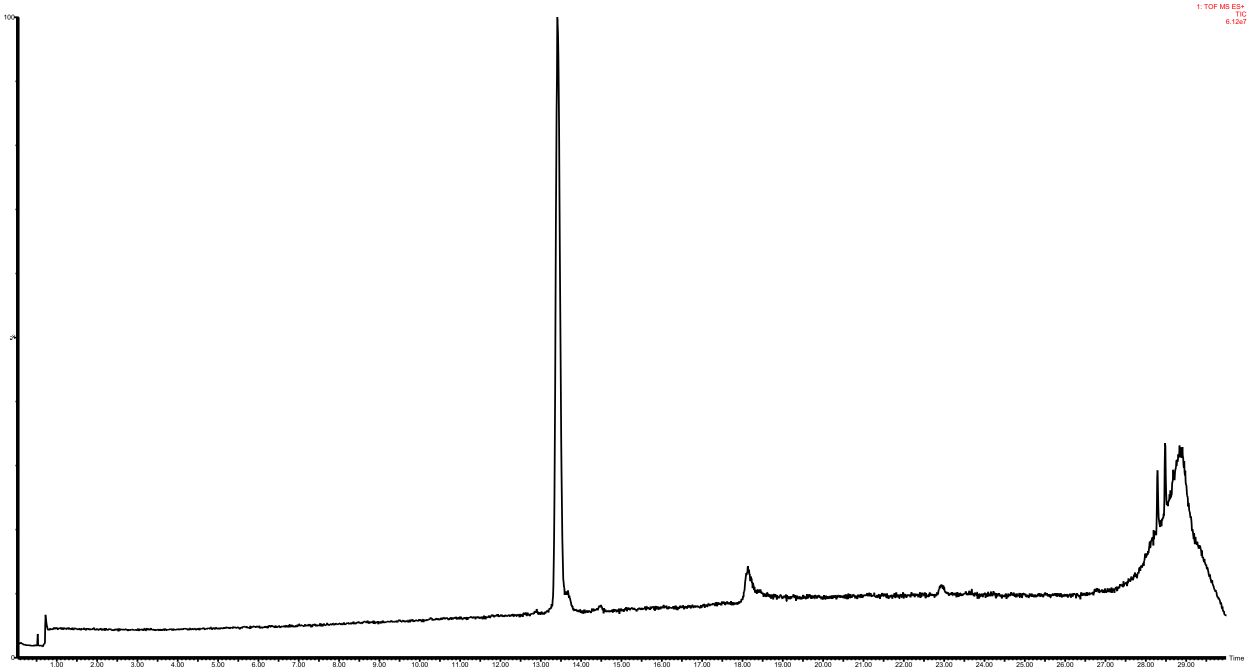


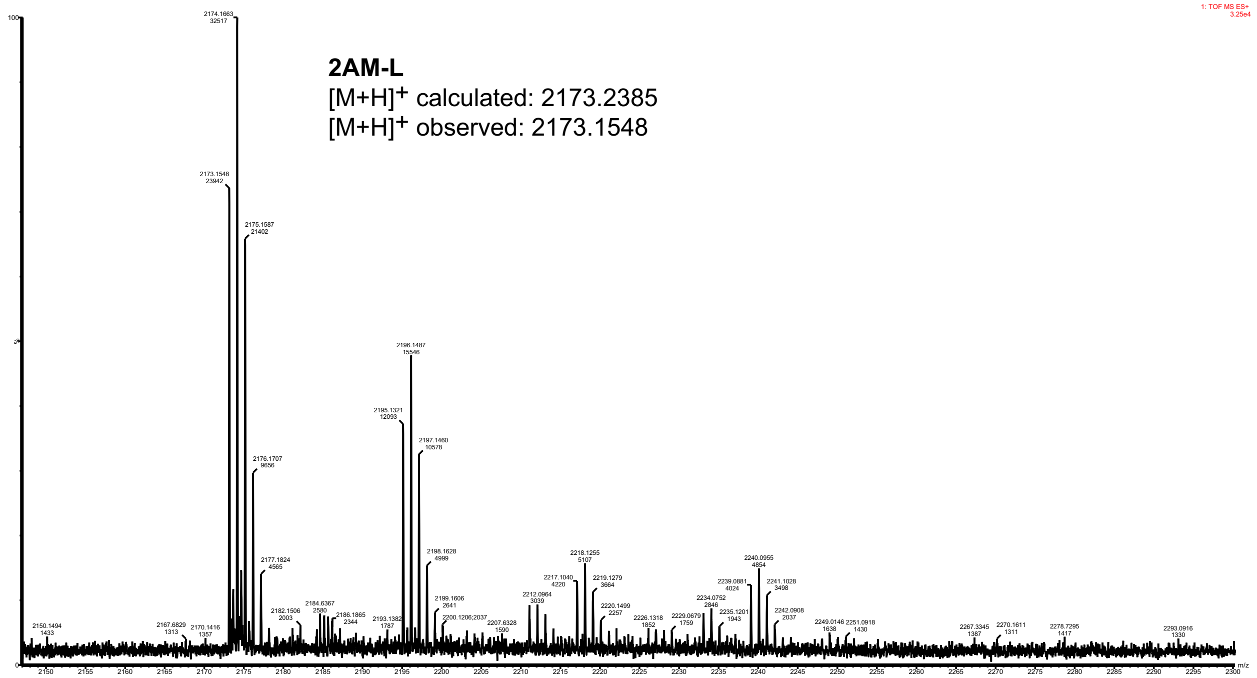

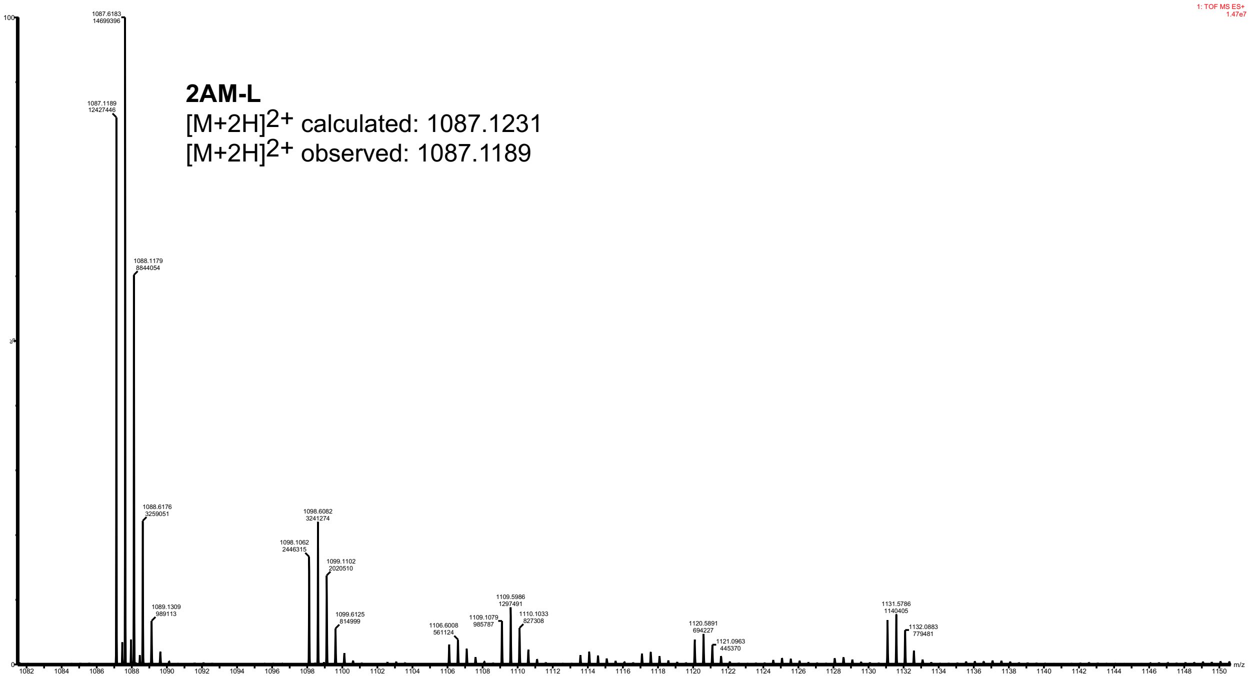
1: TOF MS ES+
1.47e7
2AM-L
[M+2H]2+ calculated: 1087.1231
[M+2H]2+ observed: 1087.1189
m/z

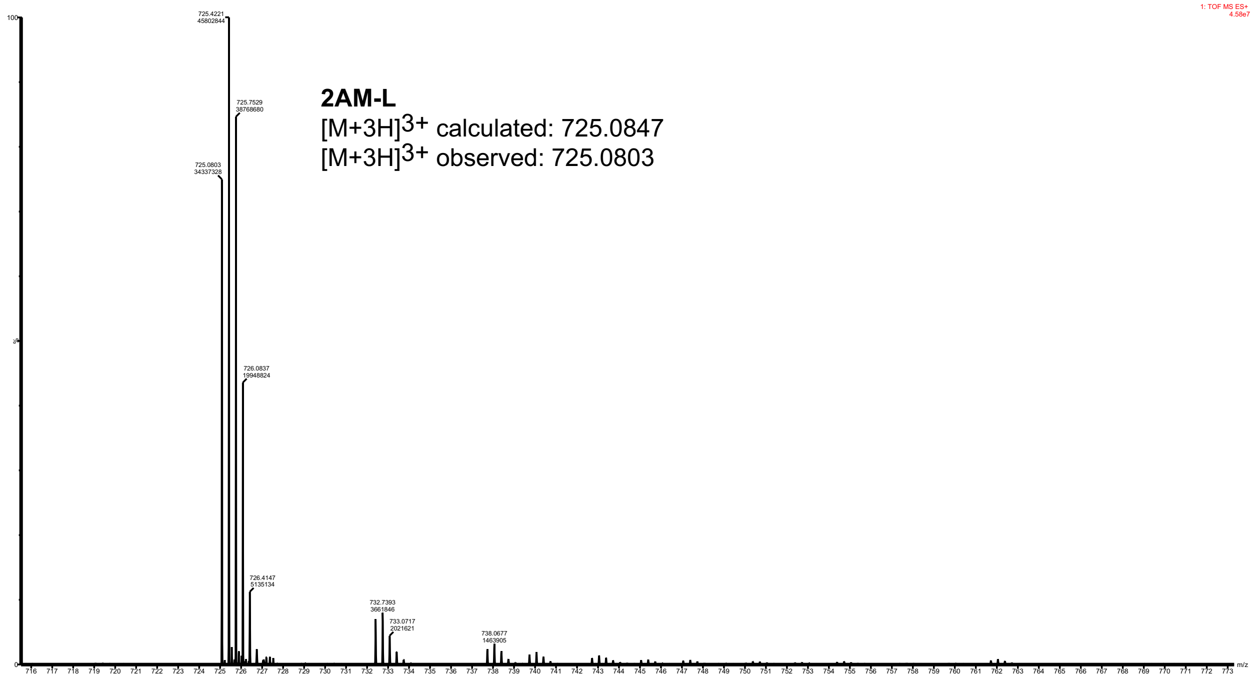
1: TOF MS ES+
4.58e7
2AM-L
[M+3H]3+ calculated: 725.0847
[M+3H]3+ observed: 725.0803
m/z

*Characterization of 2AT-L*

LC-MS of 2AT-L

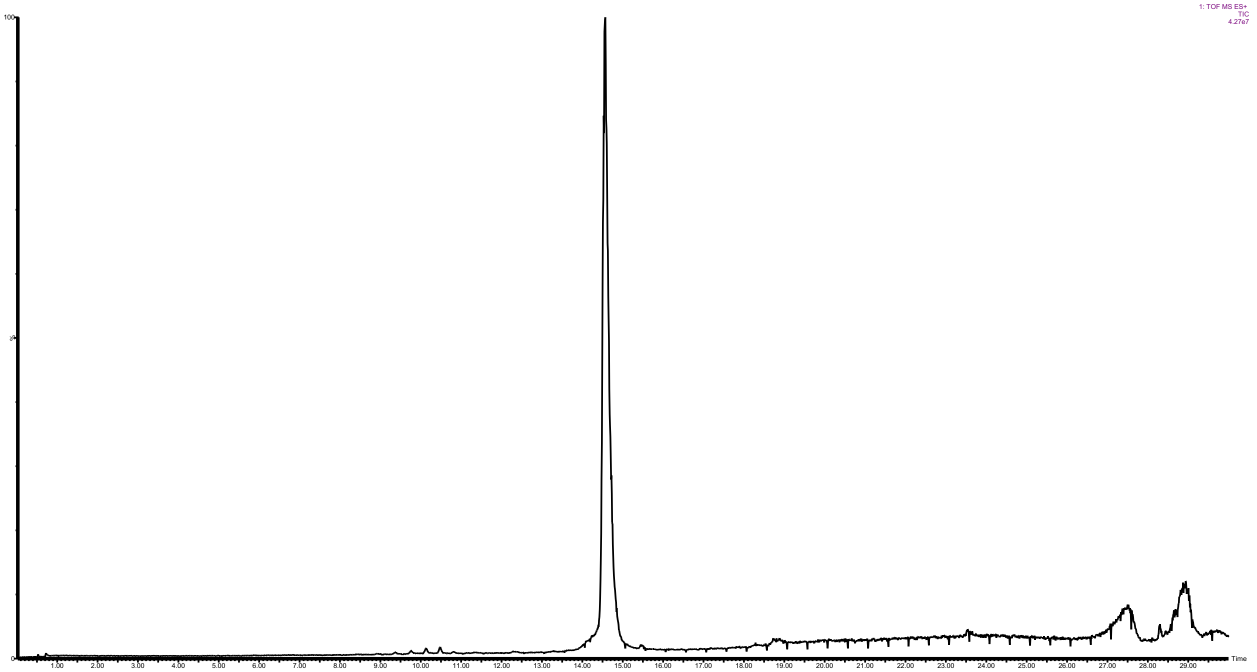

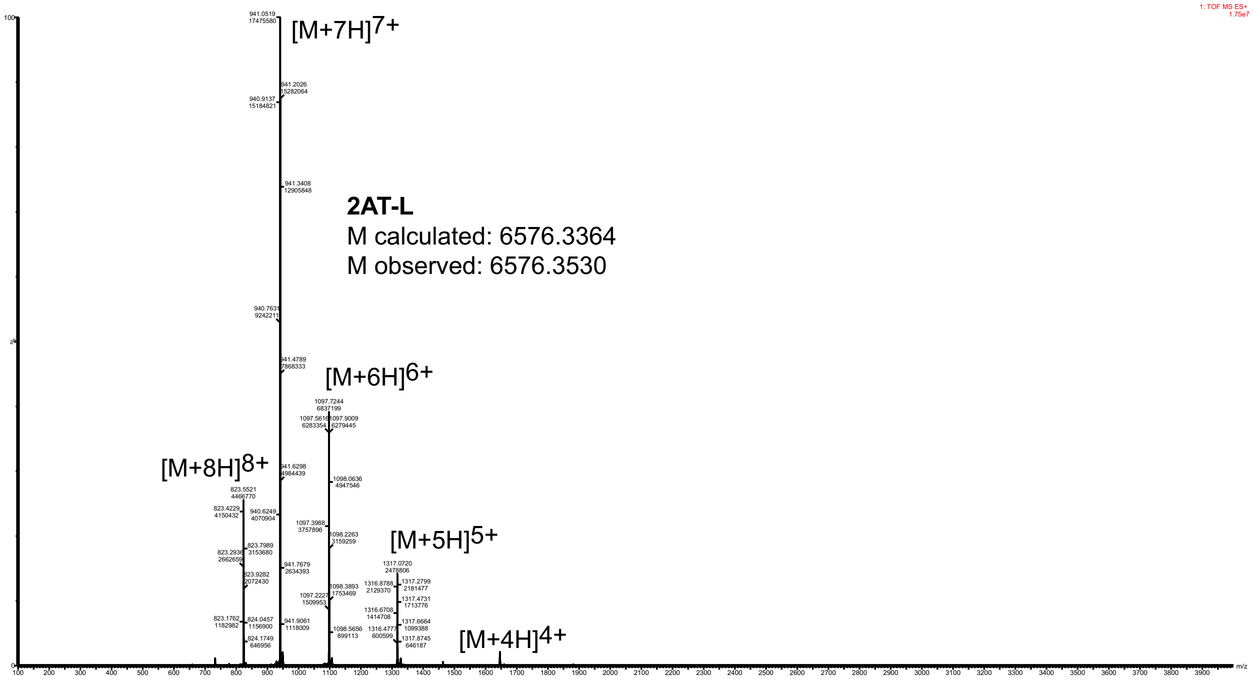

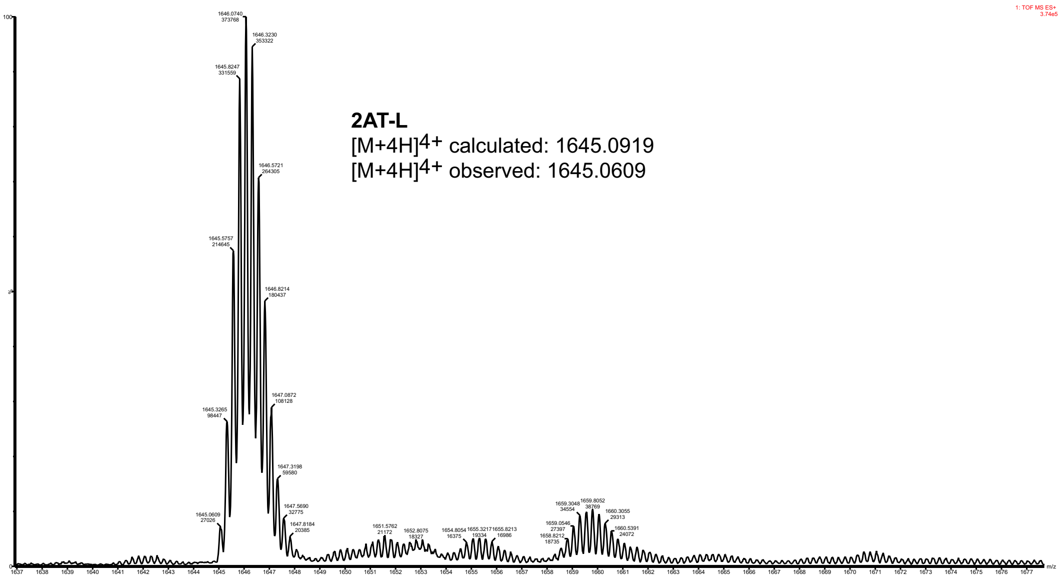
2AT-L
[M+4H]4+ calculated: 1645.0919
[M+4H]4+ observed: 1645.0609

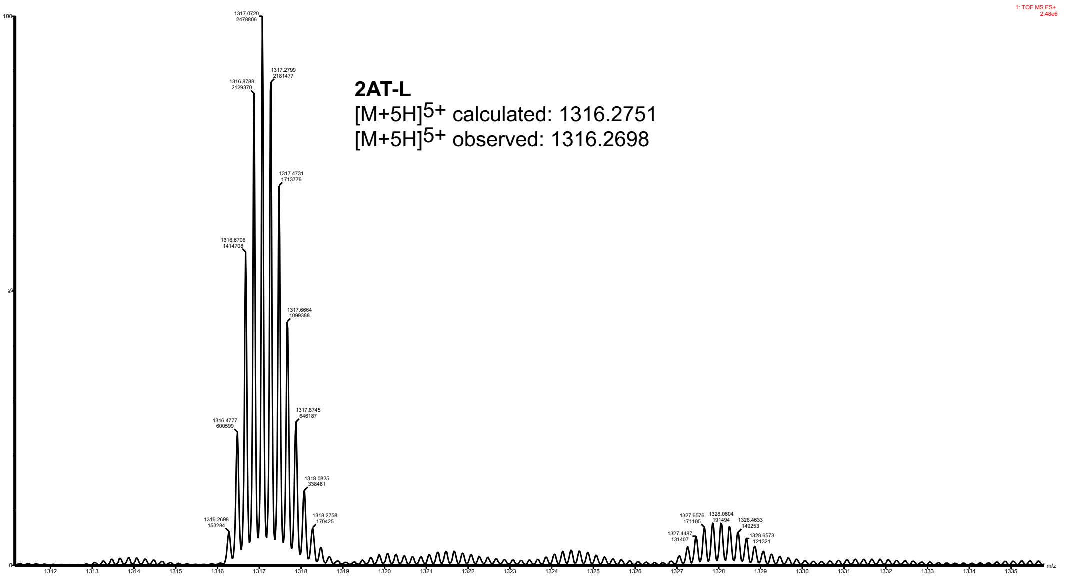
2AT-L
[M+5H]5+ calculated: 1316.2751
[M+5H]5+ observed: 1316.2698

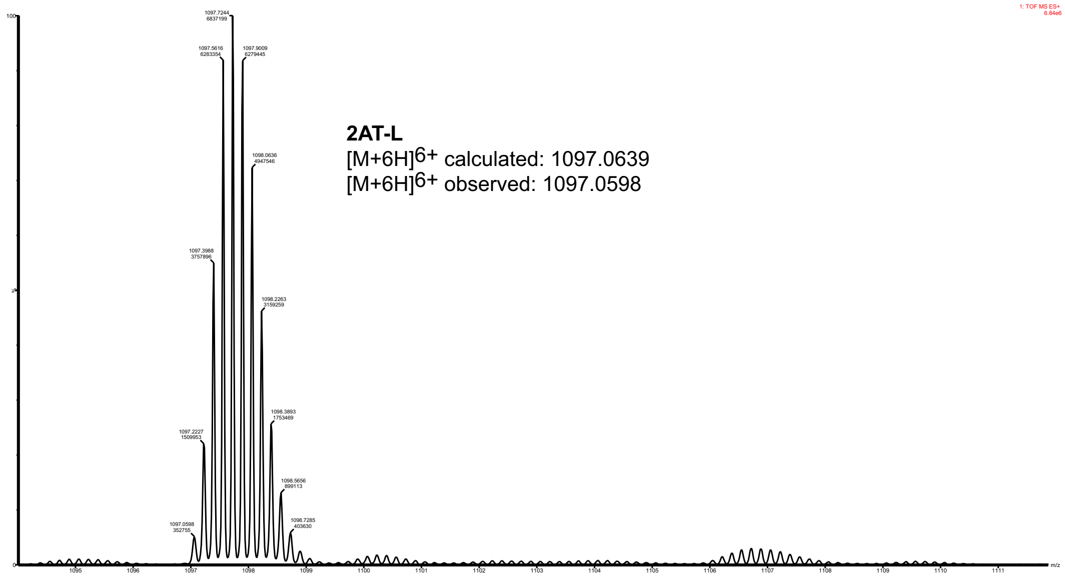
2AT-L
[M+6H]6+ calculated: 1097.0639
[M+6H]6+ observed: 1097.0598

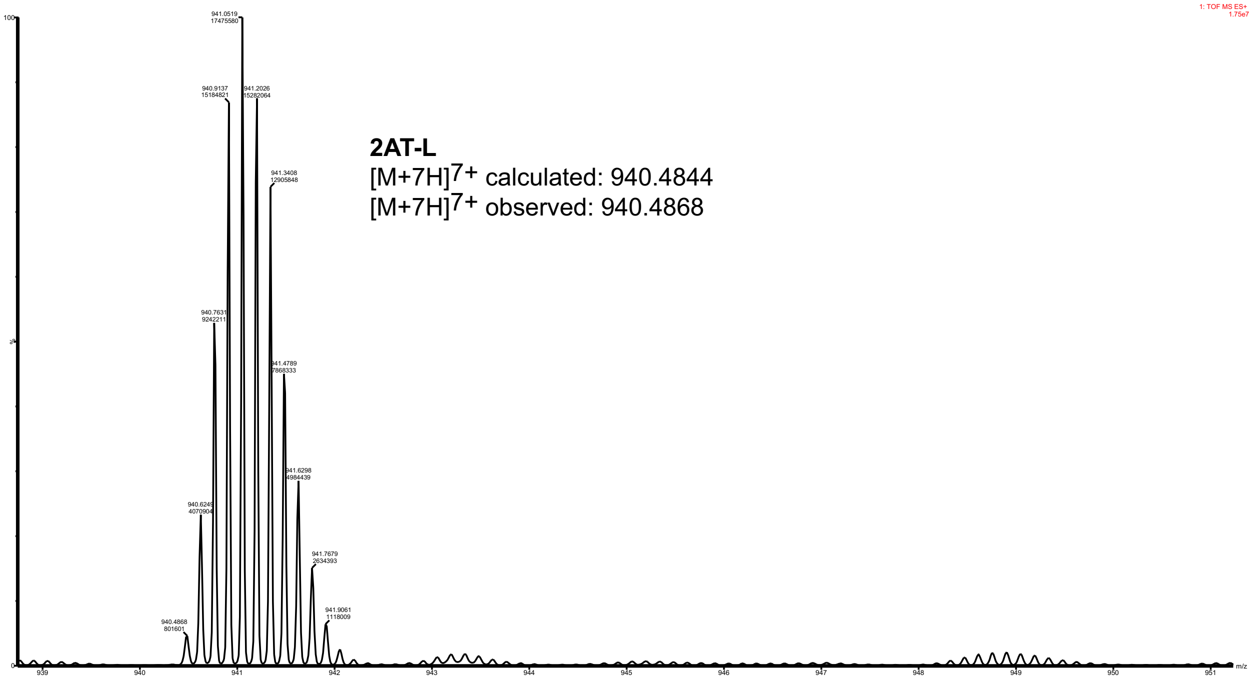
1: TOF MS ES+
1.75e7
2AT-L
[M+7H]7+ calculated: 940.4844
[M+7H]7+ observed: 940.4868
m/z

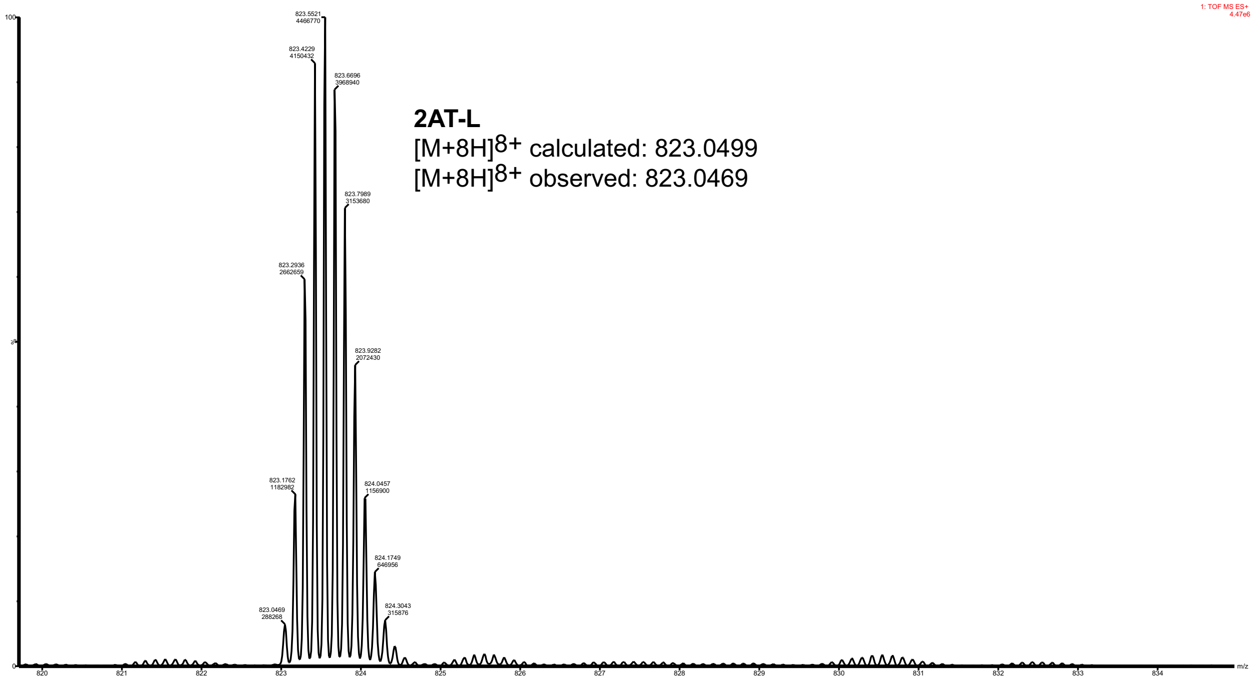
1: TOF MS ES+
4.47e6
2AT-L
[M+8H]8+ calculated: 823.0499
[M+8H]8+ observed: 823.0469
m/z
